# The *JAW-TCP-FUL* genetic axis triggers an early reorientation of cell anisotropy to initiate and drive fertilization-dependent fruit elongation in Arabidopsis

**DOI:** 10.1101/2025.05.20.655035

**Authors:** Anurag N. Sharma, Kirti, V. Avani, S. S. Nikhil Shanmugham, Ahana Jayanarayan Perumadaraya, Mariam Khadeeja, Dhanya Pradeep, Sharath Basavaraju, Kailash Chandra, Annapoorni Rangarajan, Utpal Nath

## Abstract

In angiosperms, the ovary grows and develops into a fruit after fertilization, and the seeds are formed within to ensure reproductive success. Although genetic regulators suppressing fertilization-independent fruit growth or parthenocarpy have been identified in the Brassicaceae model *Arabidopsis thaliana*, the fertilization-dependent activator of fruit growth has not been elucidated. Here, we show that the miR319-regulated TCP transcription factors (JAW-TCPs) directly activate the transcription of the *FRUITFULL* (*FUL*) gene and promote fruit morphogenesis. By activating *FUL*, JAW-TCPs indirectly repress the four valve-margin specifying genes *SHATTERPROOF1*, *SHATTERPROOF2*, *INDEHISCENT* and *ALCATRAZ* in the valves. Mutating these genes suppresses defects caused by the combined loss of *JAW-TCPs* and *FUL*. Through extensive confocal imaging studies, we deciphered the cellular basis of *JAW-TCP* function revealing that JAW-TCPs promote fruit elongation by triggering the reorientation of cell anisotropy along the length axis at an early growth stage after fertilization. Our study uncovers a fertilization-dependent genetic module driving fruit elongation and sets the stage to identify other genetic regulators in this pathway.

## Introduction

Reproductive success in angiosperms depends on successful development and functioning of their reproductive organ system - the flower^1^ - which comprises four concentric organs, namely sepals, petals, the male reproductive organ or stamens, and the female reproductive organ or pistil^2^. Mature flowers are formed within the first 13 of 20 developmental stages in the model plant *Arabidopsis thaliana*^3^. The pistil develops from stage 6 to 13 and comprises three distinct structures at maturity: a basal ovary (encasing the ovules), a central style, and a distal stigma^3^. By the end of stage 13, the flower undergoes anthesis, with the stamens aligning with the pistil such that their pollen-producing anthers lie above the stigma, enabling pollen deposition and eventually leading to fertilization of the ovules within the ovary. Subsequently, the ovules develop into seeds while the ovary concurrently matures into a fruit. As with other plant organs, mature pistil formation involves extensive cell division and moderate expansion. However, fruit elongation in Arabidopsis primarily occurs by cell expansion^4^.

Along with phytohormones which regulate flower and fruit development^5–10^, a small group of transcription factor-encoding genes are essential for pistil development and growth. At stage 6, *AGAMOUS* (*AG*) initiates the patterning of the pistil primordium^11^. Subsequently, two MADS-box genes, *SHATTERPROOF1* (*SHP1*) and *SHP2*, are expressed throughout the pistil until stage 9, when they pattern the ovules together with *SEEDSTICK* (*STK)*^12,13^. After this stage, expression of these genes becomes restricted to the sub-medial region of the ovary^12^.

The exclusion of *SHP1* and *SHP2* from the lateral region coincides with the onset of *FRUITFULL* (*FUL*) expression in the lateral ovary wall^14^ and the *REPLUMLESS* (*RPL*) expression in the replum^15^. FUL represses *SHP1* and *SHP2* transcription in the lateral ovary wall and its function extends beyond the pre-fertilization stages^16,17^. After fertilization, continued spatial separation of expression of these genes is critical for establishing the distinct domains comprising the fruit wall - the valve, valve margin, and replum^18^. In the later phase of fruit development, *SHP1* and *SHP2* activate *INDEHISCENT* (*IND*) and *ALCATRAZ* (*ALC*)^19,20^, and these four genes together specify the valve margin tissue required for fruit dehiscence and seed dispersal at maturity. Loss-of-function mutations in *FUL* result in excessive cell division in the pistil before fertilization and reduced cell expansion in the fruit valve after fertilization due to ectopic expression of valve-margin-specifying genes^14,21^. In *ful-1*, the null mutant allele of *FUL*, fruits fail to elongate, a defect that can be fully rescued at both the cellular and organ levels by mutating *SHP1*, *SHP2*, *IND* and *ALC* genes^19^. These genetic analyses underscore the importance of maintaining mutually exclusive spatial expression domains of *FUL* (in the valve) and *SHP1/2* (in the valve margin) during fruit elongation.

The transition from flower to fruit formation is a resource-intensive process requiring substantial nutritional investment^22^. Fruit formation is initiated by successful fertilization and is sustained by embryogenesis^23–25^. The fertilized egg and the developing embryo may release mobile signals to promote fruit maturation, likely by regulating genes involved in fruit morphogenesis, including *FUL* and *SHP1/SHP2*^4,14,17,21^.

Although the *FUL-SHP1/2* module is established before anthesis, it alone cannot further promote elongation of the ovary in the absence of fertilization. One explanation for this is that, different genetic modules independently establish and maintain the mutually exclusive expression domains of *FUL-SHP1/2* during pistil maturation and fruit elongation. During pistil patterning, the *FUL*-*SHP1/2* module is regulated by *JAGGED*, *FILAMENTOUS FLOWER (FIL)* and *YABBY3 (YAB3)^26^*, either directly or indirectly. Fruits of *fil;yab3* double mutants resemble wild-type fruits in length, and only patterning appears to be disrupted^26^, suggesting fruit elongation is independently regulated.

The leaf-modified origin of angiosperm flowers has long fascinated plant biologists^27^, proposing that genes responsible for the growth and development of vegetative organs would also regulate floral organs. For example, the miR319-regulated *JAW-TCPs^28^* regulate leaf morphogenesis as well as floral organs^29–32^. While miR319a is necessary to suppress *JAW-TCP* function during petal formation and stamen development^30,32^, the *JAW-TCPs* are required during pistil development to promote ovule identity and apical patterning^29,31^. Apart from their role in post-embryonic vegetative and reproductive growth, *JAW-TCPs* also promote hypocotyl cell elongation during photomorphogenesis. An interesting parallel between hypocotyl growth and fruit growth is that, both the organs grow solely by cell elongation^4^. We considered that, apart from their role during flower development^29–32^, whether *JAW-TCPs* have any role in fruit elongation.

In this study, we show that *JAW-TCPs* are essential for triggering and sustaining fruit elongation in Arabidopsis after fertilization. Our results reveal that JAW-TCPs directly activate *FUL* transcription during fruit morphogenesis while indirectly repressing valve-margin-specifying genes in the valve. The *JAW-TCP-FUL* genetic axis promotes anisotropic cell expansion in the fruit valve, contributing to directional cellular growth post-fertilization. This study provides new insights into the genetic regulation of fruit development and highlights the versatility of *JAW-TCP* function at the organ level while regulating similar cellular traits.

## Results

### JAW-TCPs are required for the elongation of fruit and valve epidermal cells

Simultaneous downregulation of five *JAW-TCPs* in the *jaw-D* mutant results in mildly altered silique architecture without changing its overall development and patterning^28^, suggesting their role in fruit growth. To determine their exact function, we generated various combinations of *JAW-TCP* loss-of-function mutants by genetic crosses and studied their fruit growth. Fruits of three single mutants (*tcp2*, *tcp4*, *tcp10*), two double mutants (*tcp2;4* & *tcp4;10*), and two triple mutants (*tcp2;4;10* & *tcp3;4;10*) were indistinguishable from Col-0 **(Fig. 1a-b, S1a**), demonstrating functional redundancy among these genes in fruit development. However, the *tcp2;4;24* triple mutant, the *tcp2;3;4;10* quadruple mutant, and the *tcp2;3;4;10;24* quintuple mutant (henceforth referred to as *tcp^Q^*) fruits displayed reduced length. In particular, *tcp^Q^*formed mature siliques >2.5-fold shorter than Col-0 (**Fig. 1a-b, S1a**), demonstrating that JAW-TCPs promote fruit elongation.

**Figure 1:**
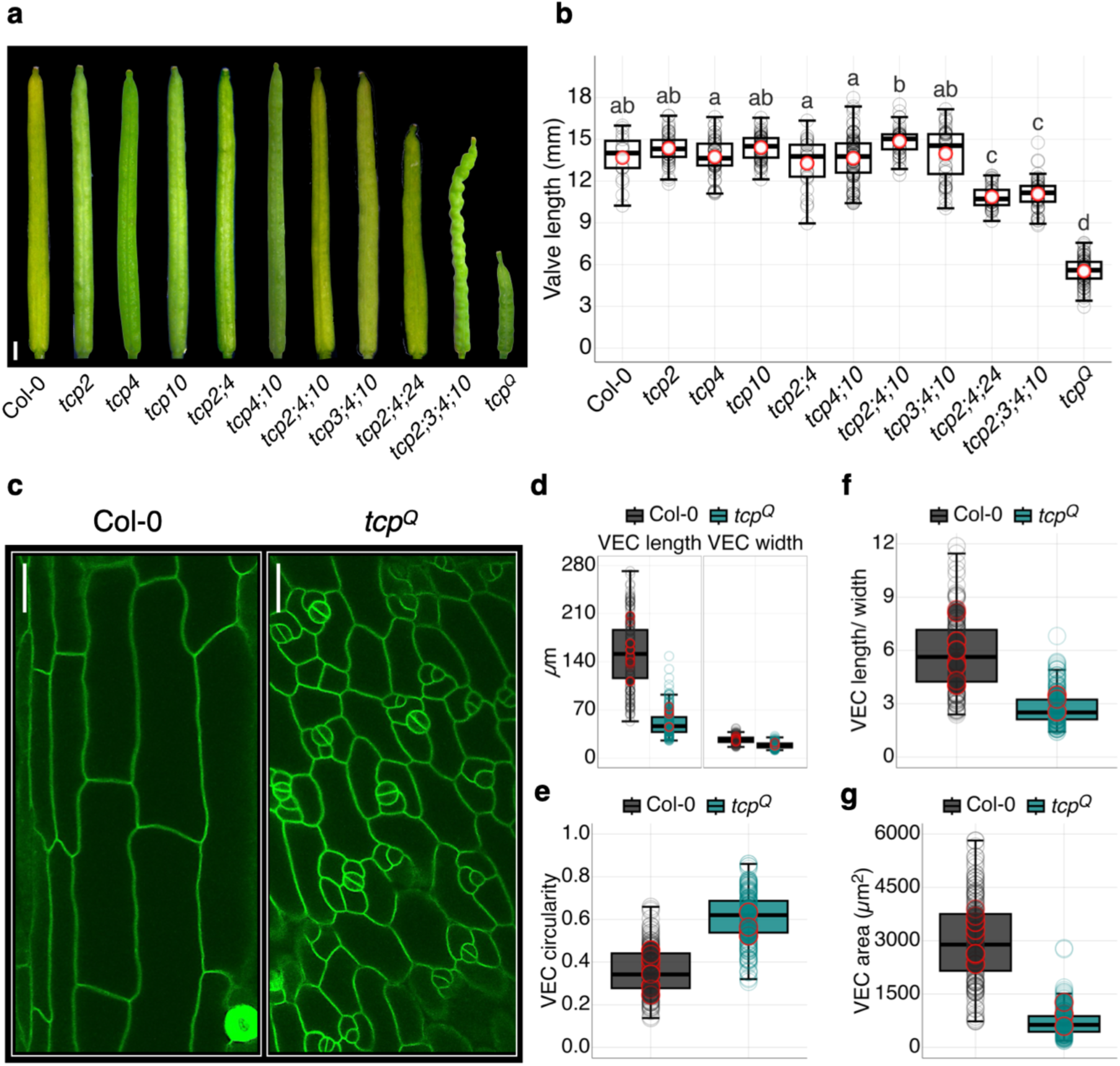
Reduced fruit length in the *jaw-tcp* loss-of-function mutants. (a-b) Images of mature fruits of indicated genotypes (a) and length of their valves (b) averaged from 20-100 fruits collected from at least three plants per batch across three independent batches. Open grey and open red circles indicate the length of individual valves and average valve length, respectively. Letters above the boxplots indicate statistical significance of *p*<0.05 as measured by a one-way ANOVA followed by Tukey’s honestly significant difference (HSD). (c) Confocal images of valve epidermal cells (VECs) of mature fruits with their cell membrane marked by *pUBQ10::ACYL-YFP*^34^ fluorescence. (d-g) Length & width (d), circularity factor (e), length/width ratio (f) and area (g) of the VECs (n = 153-300 cells) shown in (c), quantified from 3-10 fruit valves from at least three independent plants across two independent batches. Open grey/teal circles and open red circles indicate individual cell values and values averaged from each fruit. Each boxplot in (b, d-g) indicates the second and third quartile of data points, with the horizontal bar representing the median. Upper and lower whiskers indicate either the highest/ lowest value or ±1.5 times the interquartile range. Scale bars, 1 mm (a) and 20 µm (b).

The Arabidopsis fruit wall comprises two lateral valves and two medial repla, with the valve-margin tissue connecting the two along their length^33^. Earlier work showed that cell growth, not cell division, drives post-fertilization fruit elongation in Arabidopsis ^4,25^. To determine the cellular basis of reduced length in *tcp^Q^* fruits, we compared the morphology of valve epidermal cells of *tcp^Q^* and Col-0 fruits expressing the *pUBQ10::ACYL-YFP* reporter that marks the plasma membrane^34^ and reveals the cell outline. Whereas the epidermal cells in Col-0 were elongated (∼150 µm length) along the proximo-distal axis of the mature fruit, those in *tcp^Q^* were ∼3-fold shorter and more isotropic in shape with a higher circularity index value compared to the wild type (**Figures 1c-e, S1b**). However, the cell width in *tcp^Q^* remained relatively unaltered, resulting in a >2-fold reduction in the cell length-to-width ratio (**Figure 1d, 1f**). Reduced cell length in the mutant was also reflected in a more than 4-fold decrease in its average cell area (∼3000 µm^2^ in Col-0 and ∼700 µm^2^ in *tcp^Q^*) (**Figure 1g, S1b**). The overall internal tissue histology of the mutant was comparable to that of wild type, as studied in toluidine blue-stained transverse sections of stage 17-A pistils (**Figure S1c**). Together, these results provide genetic evidence that JAW-TCPs promote fruit length primarily by increasing cell elongation.

### *JAW-TCP*s and *FUL* form a genetic module in promoting fruit elongation

To determine the molecular basis of reduced fruit length in the *tcp^Q^*mutant, we estimated the transcript level of known fruit-patterning genes *FUL*, *SHP1*, *SHP2*, *IND*, *ALC* and *RPL^17,19–21^*in *tcp^Q^* gynoecia compared with their levels in Col-0 (**Figure 2a**). Similar to *ARR16*^35^, a reported direct target of JAW-TCPs, the level of *FUL* transcript was reduced in *tcp^Q^* relative to Col-0. Consequently, the activity of the *pFUL::GUS* reporter signal was completely absent from *tcp^Q^* valves, even though abundant GUS activity was observed in the Col-0 fruit (**Figure 2b**), suggesting that *JAW-TCPs* are required for *FUL* transcription. GUS activity of *pTCP4::TCP4-GUS* reporter*^31,36^* in Col-0 fruit valves was similar in pattern to that of *pFUL::GUS*, pointing to cell-autonomous activation of *FUL* transcription by JAW-TCPs (**Figures 2b, S2a-b**). Corroborating this finding, expression of the *pTCP4::FUL* transgene rescued the fruit length defect in the *ful-1* mutant background to a large extent (**Figures S2c-d**), suggesting that *JAW-TCP* expression in the fruit valves is required for *FUL* transcription and *FUL*-mediated fruit elongation. The *pTCP4::TCP4-GUS* reporter activity was not altered during the early growth stages in *ful-6* (a hypomorphic allele of *FUL*) fruits^16^ (**Figure S2a**), suggesting that *JAW-TCP* expression is not dependent on *FUL*.

**Figure 2.**
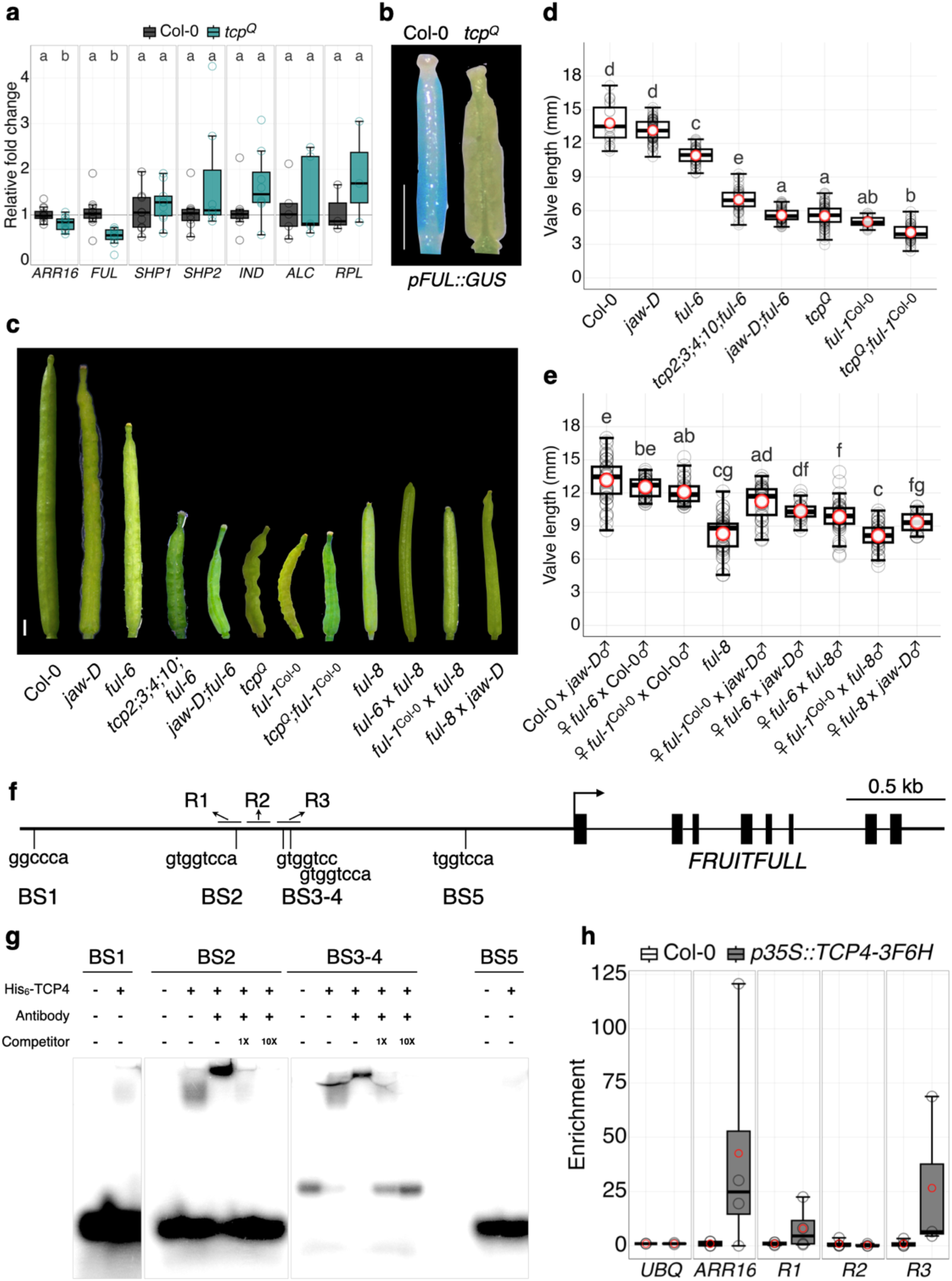
Reduced expression and activity of *FUL* in the absence of *JAW-TCP*s. (a) Relative transcript level of genes involved in fruit elongation as determined by RT-qPCR analysis from the total RNA isolated from stage 14-15 gynoecia of Col-0 and *tcp^Q^*. Each open circle indicates the value of one biological replicate. The letters above the boxplots indicate a statistical significance of *p*<0.05, as determined by a Welch t-test. (b) GUS reporter analysis of Col-0 and *tcp^Q^*stage 16 fruits expressing *pFUL::GUS*. (c-e) Images of mature fruits of the indicated genotypes (c) and length of their valves (d, e). Averages (red circles) of 23-81 (d) or 18-45 fruits (e) are shown. Fruits were collected from at least three plants per batch across three independent batches. Open grey circles indicate the length of individual valves. The difference in letters above the boxplots (d-e) between genotypes indicates the statistical significance of *p*<0.05 as measured by a one-way ANOVA followed by Tukey’s HSD. (f) A schematic representation of the *FUL* genomic region marking five putative TCP4-consensus binding sites (BS1 to BS5) in the upstream regulatory region. R1-R3 represent the sites tested for TCP4-binding in the ChIP-qPCR assay in (h) below. Filled vertical boxes represent exons. (g) Image of an autoradiogram of electrophoretic mobility shift assay analysis showing the binding of His_6_-TCP4 recombinant protein with the indicated P^32^-radiolabelled oligonucleotides corresponding to BS1-BS5 shown in (f) above. + and - indicate the presence and absence of indicated compounds, respectively. α-His_6_ antibody and the corresponding non-radiolabeled oligonucleotides were used for the super-shift assay and the competition assay, respectively. (h) ChIP-qPCR assay testing the in vivo binding of the TCP4-3F6H protein at the R1-R3 binding sites indicated in (f). *UBIQUITIN10* (*UBQ*) and *ARABIDOPSIS RESPONSE REGULATOR 16^35^* (*ARR16*) were used as negative and positive controls, respectively. Each boxplot in (a, d, e, h) indicates the second and third quartile of data points, with the horizontal bar representing the median. The upper and the lower whiskers indicate either the highest/ lowest value or ±1.5 times the interquartile range. Scale bars, 1 mm (b, c).

To genetically validate the *JAW-TCP-FUL* gene expression studies described above, we combined incomplete loss of function alleles of *JAW-TCP*s (*jaw-D* or *tcp2;3;4;10*) and of *FUL* (*ful-6*) and studied their fruit phenotype. In both *jaw-D;ful-6* and *tcp2;3;4;10;ful-6* mutants, fruits were shorter compared to their respective parents (**Figure 2c, 2d, S2e**), though the *jaw-D;ful-6* fruit transverse section appeared similar to Col-0 (**Figure S2f)**. This result can be interpreted as *JAW-TCP*s and *FUL* working in the same molecular pathway, although the possibility of them working in a parallel pathway cannot be ruled out. To test the latter possibility, we generated the *tcp^Q^*;*ful-1*^Col-0^ hextuple mutant by combining the null alleles of *JAW-TCP*s and *FUL,* which produced fruits of length similar to their parental lines (**Figure 2c, 2d**), providing more credence to the conclusion that *JAW-TCP*s and *FUL* work in the same molecular pathway with *FUL* being dependent on JAW-TCPs for its transcriptional activity during fruit growth.

### JAW-TCPs directly activate *FUL* transcription

The *JAW-TCP* genes encode DNA-binding transcription factors whose consensus binding sites have been well characterized^37^. We analysed the upstream regulatory sequences of the *FUL* locus for the presence of JAW-TCP binding sites and identified five putative sequence motifs where these proteins can potentially bind (BS1-5; **Figure 2f**). In an electrophoretic mobility shift assay, a recombinant TCP4-His_6_ fusion protein retarded the mobility of ^32^P-labelled synthetic oligonucleotides corresponding to BS2 and BS3-4 but not to BS1 and BS5 (**Figure 2g**). The mobility of these DNA-protein complexes was further retarded by the inclusion of an α-His_6_ antibody in the binding reaction, suggesting that the first retardation was indeed due to the TCP4-His_6_ protein. Adding nonradioactive oligos to the reaction mixture abolished the autoradiogram signal of the DNA-oligo complex, indicating that the binding of TCP4-His_6_ to the oligos is specific. Further, a recombinant TCP4-3F6H protein was found to be recruited to the *FUL* upstream regulatory regions corresponding to BS2 and BS3-4 (R1 & R3 regions in **Figure 2h**) in a chromatin immunoprecipitation (ChIP) experiment followed by qPCR using α-Flag antibody and total chromatin isolated from the *35S::TCP4-3F6H* transgenic line^38^. No enrichment was observed in the area corresponding to R2, the intervening region between BS2 and BS3-4. These results suggest that JAW-TCPs bind to the *FUL* locus at BS2 and BS3-4 sites and activate its transcription.

To further test the importance of BS3-4 in JAW-TCP-mediated *FUL* activation, we generated Arabidopsis lines with mutated BS3-4 sequence using CRISPR/Cas9 editing. We established a Cas9-free stable homozygous line (henceforth called *ful-8*) where the GTGG**T**CC cis-element was edited to GTGG**G**CC (**Figure 2e, S2e, S2g**) with a T→G transversion (bold-faced). Morphometric analysis revealed that *ful-8* produced mature fruits shorter than Col-0 fruits by ∼40% (**Figure 2c, 2e**), an extent of phenotypic change intermediate between *ful-6* (a hypomorphic allele) and *ful-1* (a null allele). Fruit length in the F1 individuals of the genetic crosses of known *FUL* alleles such as *ful-6* or *ful-1*^Col-0^ with *ful-8* (*ful-6*♀ x *ful-8*♂ and *ful-1*^Col-0^♀ x *ful-8*♂ genotypes, respectively) produced fruits shorter than those resulting from their corresponding crosses with wild type (*ful-6*♀ x Col-0♂ or *ful-1*^Col-0^♀ x Col-0♂ genotypes, respectively) (**Figures 2c, 2e, S2e**), providing genetic evidence that *ful-8* is indeed a new loss-of-function allele of *FUL* and its shorter fruit phenotype is perhaps not due to mutations elsewhere in the genome. Together with earlier results, we concluded that JAW-TCPs directly activate *FUL* transcription by binding to its upstream regulatory sequences and promote fruit elongation.

### The rate of elongation during early fruit growth is reduced in the *jaw-tcp* mutants

Under standard growth conditions, a typical wild-type pistil is fertilized at stage 14, breaking the temporary growth pause that occurs after pistil maturation. Fertilization triggers fruit elongation through stages 15 to 17-A, eventually turning yellow at the end of stage 17-B, marking fruit maturity^3,18^. To determine how the growth of *jaw-tcp* loss-of-function fruits deviates from that of wild-type, we analyzed the growth kinematics of *tcp^Q^*and *jaw-D;ful-6* fruits across stages of maturation and at increasing days after pollination (DAP) to saturation and compared the values with those for Col-0 and *ful-1*^Col-0^ fruits^4^ (**Figure 3**). On an approximation, stages 14-15 correspond to 0-2 DAP, whereas stage 16 continues for another day, leaving the remaining ∼7-8 days of fruit growth for stage 17, wherein most elongation occurs^39^.

**Figure 3.**
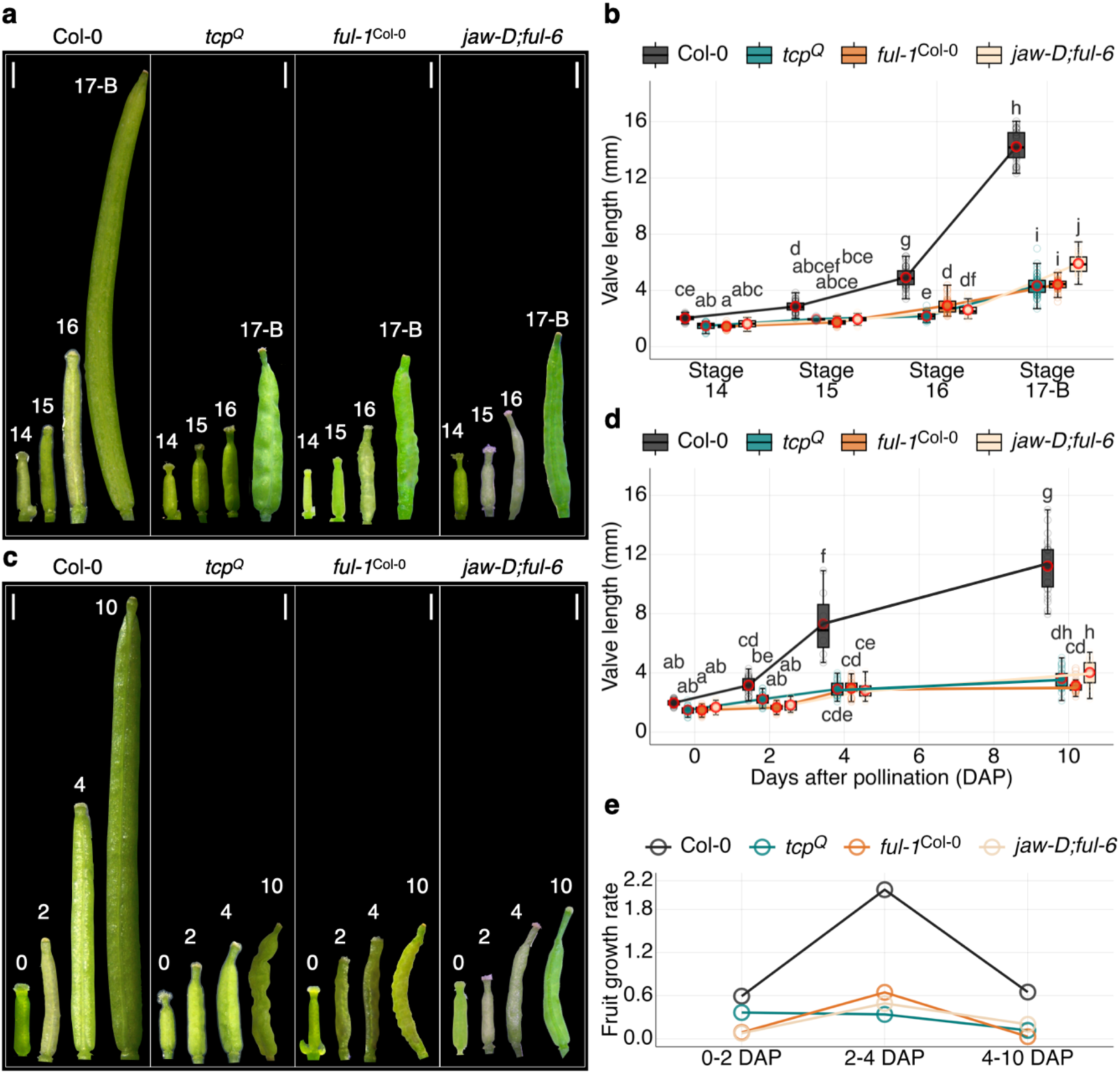
Kinematic analysis of fruit elongation in the mutants of *JAW-TCP*s and *FUL*. (a, b) Images of stage 14 to stage 17-B fruits of the indicated genotypes (a) and their valve lengths (b). The staging was done as previously reported^3^. (c, d) Images of fruits at 0, 2, 4 and 10 days after pollination (DAP) (c) and their valve lengths (d). (e) Rate of valve elongation of the indicated genotypes as derived from data shown in (d). Data in (b) and (d) are from 21-58 and 18-33 fruits, respectively. Fruits were collected from at least three plants per batch across three independent batches. Open colored and open red-coloured circles indicate the length of the individual valves and the average valve length, respectively. The difference in letters above the boxplots in (b, d) between genotypes indicates a statistical significance of *p*<0.05 as measured by a two-way ANOVA followed by Tukey’s HSD. Each boxplot in (b) and (d) indicates the second and third quartile of data points, with the horizontal bar representing the median. The upper and the lower whiskers indicate either the highest/ lowest values or ±1.5 times the interquartile range. Scale bars, 1 mm (a, c).

Col-0 fruits nearly doubled from stage 14 to stage 15 and continued to elongate through stage 16 until their maturity at 17-B, finally measuring ∼14 mm long (**Figures 1a-b, 2c-d, 3a-b and S3a-b**). Although a similar trend in elongation pattern was observed for *tcp^Q^*, *ful-1*^Col-0^ and *jaw-D;ful-6* fruits, the extent of growth was much less in these mutants (**Figure 3a-b, S3a-b**), resulting in the final fruit length of 5-6 mm at stage 17-B (**Figure 2c-d, 3a-b)**. To study the kinematics of fruit growth, we compared fruit elongation in all four genotypes at 0, 2, 4 and 10 DAP (**Figure 3c-d, S3c**). In Col-0 fruits, growth followed a typical sigmoidal kinetics (**Figure 3d**) with the rate of growth ∼0.6 mm/ day in the first 2 DAP, which sharply increased to a maximum value of ∼2 mm/ day from 2-4 DAP, before decreasing to nearly the initial value from 4-10 DAP (**Figure 3e and S3d**). Thus, the fastest relative growth occurred within the first 4 days after pollination out of a total of 10-11 days of fruit elongation. Although the initial rate of elongation of mutant fruits in the first two DAP was comparable to that of the wild type, the rate in 2-4 DAP was 0.4 mm/ day, nearly 5-fold less than that of Col-0. The rate decreased slowly but steadily thereafter to a value of ∼0.2 mm/ day from 4-10 DAP. At 10 DAP, the fruits in all four genotypes attained nearly 80% of their respective length at stage 17-B (compare **Figure 3c-d** with **Figure 3a-b, S3e**), suggesting that, though the duration of fruit growth remains unaltered, the rate of elongation is reduced in the mutant fruits, notably within the initial few days after pollination.

### An early reorientation in the anisotropy of the valve epidermal cells triggers fruit elongation

To uncover the cellular basis underlying the difference in growth between the wild type and the mutant fruits, we analyzed the growth parameters of their valve epidermal cells at 0, 2, 4 and 10 DAP (**Figure 4a-d, S4a**). At 0 DAP, epidermal cells in Col-0, *tcp^Q^* and *jaw-D;ful-6* were comparable in their surface area (100-175 µm^2^) and more isotropic in shape with length-to-width ratio ranging from 1.7 to 1.8 and circularity parameter from 0.65 to 0.73 (**Figure 4a-d**). The cell area in Col-0 increased ∼5-fold between 0-2 DAP, primarily due to growth in length, as the length-to-width ratio increased ∼2-fold during this period (**Figure S4a**). Consequently, cell anisotropy increased with the circularity parameter reducing from 0.65 to 0.45 (**Figure 4d**). Preferential growth along the length axis continued from 2-10 DAP, with the length-to-width ratio increasing ∼2-fold at 10 DAP. The circularity value decreased only marginally to 0.40 in this 8-day growth period. Thus, the cellular growth anisotropy during Col-0 fruit elongation is rapidly established within the first two days after pollination and changed to a small extent thereafter, even though cell area increased ∼3-fold from 2-10 DAP (**Figure 4b**).

**Figure 4.**
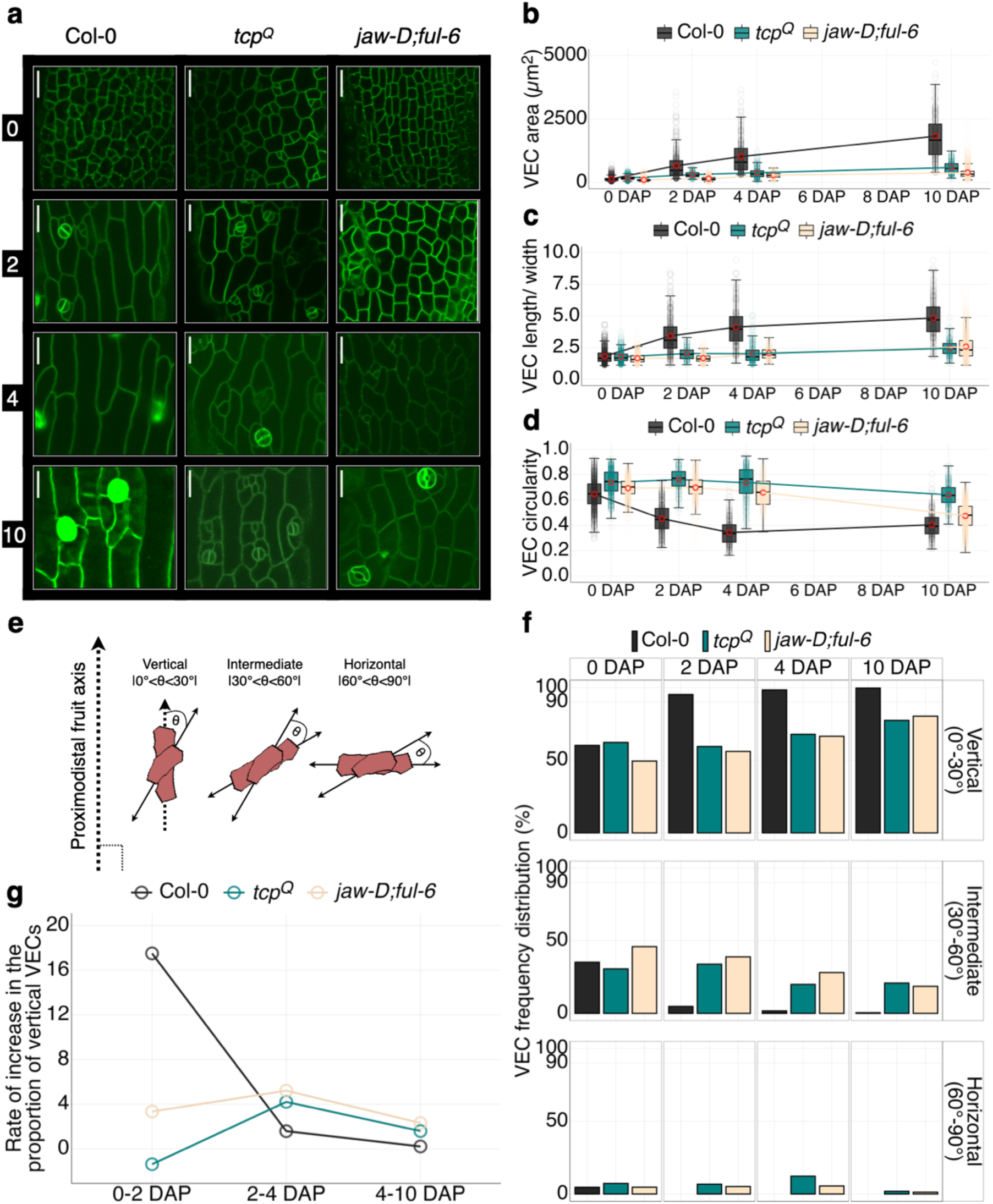
Vertical reorientation of valve epidermal cells during fruit elongation. (a) Confocal fluorescence images of mature valve epidermal cells (VECs) in the indicated genotypes (on the top) marked with *pUBQ10::ACYL-YFP* at indicated days after pollination (on the left). Scale bar, 20 µm. (b) Area, (c) length/ width ratio (d), and circularity values (d) of the VECs shown in (a). (e) A schematic showing an empirical classification of VECs into vertical, intermediate and horizontal categories based on the angle between their length axis and the proximodistal axis of the fruit at 0 DAP. (f) Frequency distribution of the VECs analyzed in (b-d) among vertical, intermediate and horizontal categories in the indicated genotypes at 0, 2, 4 & 10 days after pollination (DAP). (g) The rate of increase in the percentage of vertical cells shown in (f) between the indicated days of growth shown along the X-axis. Values in (b-d, f) were averaged from 3-10 valves from at least three independent plants across two independent batches of plants. Individual means are plotted as open red circles in (b-d). Filled circles in other colours indicate individual VEC parameter values. Each boxplot in (b-d) indicates the second and third quartile of data points, with the horizontal bar representing the median. The upper and the lower whiskers indicate either the highest and lowest value or ±1.5 times the interquartile range.

In contrast to the wild type, only a ∼1.5-fold increase in epidermal cell area was observed in *tcp^Q^* and *jaw-D;ful-6* fruits between 0-2 DAP, and the growth was ∼2.5-fold in the next eight days, similar to Col-0 (**Figure 4a, 4b, S4a**). Notably, the lack of growth in the mutant cells was observed preferentially along the length axis, with a barely detectable change in the length-to-width ratio or in the circularity value in the first two days of fruit growth. Between 2-10 DAP, although the cell area and cell length increased by ∼2.3-fold and ∼1.8-fold, respectively, the cell anisotropy value dropped only marginally, especially in *tcp^Q^* fruits. These results suggest that the *JAW-TCP-FUL* genetic axis promotes an initial and rapid increase in cell anisotropy, which underlies wild-type fruit elongation.

As mentioned earlier, fruit elongation in Arabidopsis is driven entirely by cell elongation^4,25^, as cell number remained unaltered in Col-0, *tcp^Q^* and *jaw-D;ful-6* throughout the growth period (**Figure S4b**). To determine the direction of anisotropy during the growth of valve epidermal cells described in the previous section, we measured the angle (θ) between the length axis of individual cells and the proximodistal axis of the fruit. Based on the θ-value, we empirically grouped the valve epidermal cells into three categories - vertical (|0°<θ<30°|), intermediate (|30°<θ<60°|) and horizontal (|60°<θ<90°|) (**Figure 4e**). Frequency distribution analysis of the cell categories revealed that nearly half of the cells were oriented vertically in Col-0, *tcp^Q^* and *jaw-D;ful-6* fruits at 0 DAP, whereas 30-40% cells were of an intermediate type and a small fraction of cells (∼5%) were oriented horizontally (**Figure 4f**). In Col-0, the proportion of vertical cells increased to ∼95% within the first two days of growth, with a concomitant decrease in the proportion of cells in the other two categories to negligible levels (**Figure 4g**). At 10 DAP, nearly all cells in Col-0 (>99.5%) were oriented vertically. In stark contrast to Col-0, the proportion of vertical cells remained unaltered in *tcp^Q^* and *jaw-D;ful-6* fruits at 2 DAP, with nearly 50% remaining in intermediate and horizontal categories (**Figure 4f-g**). From 4-10 DAP, the vertical cells increased in proportion to a smaller extent than the wild type, reaching ∼80% at 10 DAP, with the remainder of cells oriented in the intermediate and horizontal directions.

In the experiments described above, we considered the possibility of the confounding effect of emasculation and artificial pollination on the fruit growth analysis, especially since genetic evidence indicates that the live organs in the floral whorls surrounding the fruit inhibit its growth^24,40^. Therefore, we repeated the cell growth study at various stages in the fruits resulting from natural pollination. The overall pattern of cellular growth from stage 14 to stage 17-B corresponded with those observed in the kinematic experiments for all genotypes (compare **Figure 4** with **Figure S4c-h**). The epidermal cell area of Col-0 increased from ∼100 µm^2^ at stage 14 to ∼450 µm^2^ at stage 16, thus marking a ∼4.5-fold increase in size at an early growth phase before reaching a final size of ∼3000 µm^2^ (**Figure S4c, d**). The cell anisotropy also increased correspondingly across the stages (**Figure S4e, f, g**), notably early in growth from stages 14-16, during which a maximum decrease in the circularity index was achieved (from ∼0.65 to ∼0.4). At stage 14, only about half of the epidermal cell population was vertically oriented, the proportion of which sharply increased to a near maximum value of ∼98% by stage 16 (**Figure S4h**). The size and shape parameters of the epidermal cells in the *tcp^Q^*, *ful-1^Col-0^*and *jaw-D;ful-6* valves were similar to wildtype values at stage 14 but, in contrast to Col-0, changed little across the later growth stages (**Figure S4c-g**). The cell area in the mutant valves increased from ∼80-180 µm^2^ at stage 14 to ∼250-650 µm^2^ at stage 17-B, and the length-to-width ratio and circularity parameter remained nearly unaltered. Consequently, the proportion of vertical epidermal cells in these mutant valves increased to a much lesser extent than that in the wild type during fruit elongation, leaving a large fraction of cells in intermediate and horizontal orientation even in the mature fruit at stage 17-B (**Figure S4h**). Taken together, these data suggest that *JAW-TCPs* are required for reorienting valve cell anisotropy along the growth direction of the fruit at an early stage.

### The *JAW-TCP-FUL* genetic axis promotes fruit length by suppressing the valve margin genes

The shorter-fruit phenotype of *ful-1* is progressively rescued with the inactivation of an increasing number of the four valve margin-specifying genes, namely *SHP1*, *SHP2*, *IND* and *ALC*^19^, suggesting that FUL promotes fruit elongation by repressing these genes within the valves, thus restricting them to the margin layers in wild type^14^. These margin genes are ectopically expressed in the mesodermal cells of the *ful-1* valve, leading to increased lignification at a later stage of fruit development. *SHP2* was also ectopically expressed in the *tcp^Q^*valves as abundant GUS signal was detected throughout the valves of the *tcp^Q^*;*pSHP2::GUS* fruit (**Figure 5a-b**). Increased lignification was observed in the *jaw-D;ful-6* valves (**Figure S5a**), further suggesting an ectopic presence of the valve margin genes. To test whether the shorter-fruit phenotype in the *jaw-tcp* mutants is due to this ectopic expression, we simultaneously targeted *SHP1*, *SHP2*, *IND* and *ALC* in Col-0, in *ful-1^Col-0^* and in *jaw-D;ful-6* lines by CRISPR/Cas9 genome editing. In several T1 transformants (10 independent lines) in the *ful-1^Col-0^* background, the short-fruit phenotype of *ful-1* was rescued to a length observed in the corresponding CRISPR-edited plants (20 independent lines) in the Col-0 background (**Figure 5c-d**), as was demonstrated earlier by genetic interaction studies^19^. Strikingly, five independent T1 plants in the *jaw-D;ful-6* background also formed fruits where the short-fruit phenotype was rescued nearly to the level of the fruits where the valve-margin genes were mutated in the Col-0 background. In addition, these fruits with edited valve-margin genes were indehiscent at maturity (**Figure 5c-d**), confirming that the target genes were indeed inactivated in these lines. Supporting this observation, we also noted that the fruits edited for the target-genes showed reduced lignification compared to their unedited counterparts (**Figure S5a**).

**Figure 5.**
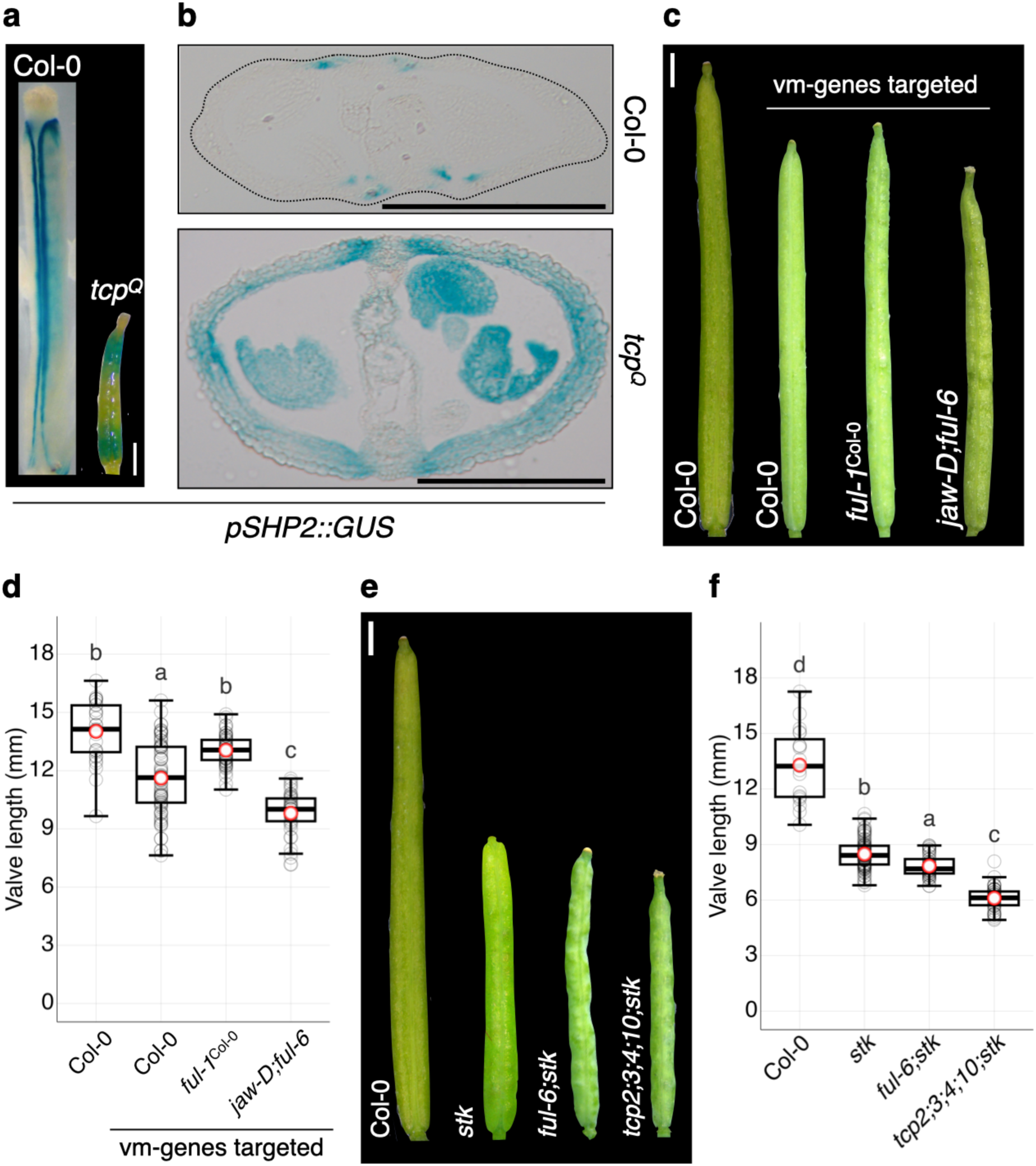
Genetic interaction between *jaw-tcp* and other mutants with altered fruit length. (a, b) *pSHP2::GUS* reporter analysis in Col-0 and *tcp^Q^* fruits at stage 17-A and the transverse sections (b) of the fruits shown in (a). (c-f) Images of mature fruits of the indicated genotypes (c, e) and their respective valve length (d, f) averaged from 29-88 fruits. ‘vm-genes targeted’ in (c-d) refers to transgenic plants expressing a Cas9-editing construct targeting a single gRNA per gene to each of the four vm-genes – *SHP1*, *SHP2*, *IND* and *ALC*. Fruits were collected from at least three plants per grown batch across three independent batches. The difference in letters above the boxplots in (d, f) between genotypes indicates the statistical significance of *p*<0.05 as measured by a one-way ANOVA followed by Tukey’s HSD. Open grey and open red circles in (d, f) indicate the individual fruit valve length and average valve length, respectively. Each boxplot in (d) and (f) indicates the second and the third quartile of data points, with horizontal bars representing the median. The upper and lower whiskers indicate either the highest/ lowest value or ±1.5 times the interquartile range. Scale bar, 200 µm (b), 1 mm (a, c, e).

While studying the *TCP4* promoter activity, we noticed the presence of TCP4-GUS signal in the entire valve region including the valve margin of the Col-0;*pTCP4::TCP4-GUS* fruit (**Figure S2b**), raising the possibility that JAW-TCPs are active in the margin cells as well, a tissue from where *FUL* is absent^21^. To test any possible role of *JAW-TCPs* in the valve margin, we expressed *MIR319A* under the regulation of the *SHP2* promoter to reduce the level of *JAW-TCPs* in this region. The resultant *pSHP2::MIR319A* fruits were indistinguishable from Col-0 fruits in length and indehiscence property (**Figure S5b-c**), suggesting that JAW-TCPs are inactive in the valve margin. The TCP4 protein did not physically interact with any valve margin-specifying proteins in a yeast two-hybrid assay (**Figure S5d**), further suggesting that JAW-TCPs are unlikely to regulate these proteins at the post-translational level. Together, these results provide genetic evidence that the *JAW-TCP-FUL* module promotes fruit elongation by suppressing the valve margin genes within the valves.

### JAW-TCPs promote fruit elongation in parallel to STK

Genetic interaction studies described above (**Figure 2c, d**) showed a small but significant decrease in the length of *tcp^Q^*;*ful-1^Col-0^*fruits compared to *tcp^Q^*. This result can be interpreted as either of or both the parental lines being not completely null, or JAW-TCPs promoting fruit elongation to a small extent in a *FUL*-independent manner which is revealed in the *ful-1*^Col-0^ background. To test the latter possibility, we generated a transgenic line expressing *pFUL::rTCP4-GR* in the *ful-1^Col-0^* background and analysed its fruit length. Whereas the length of the *ful-1^Col-0^;pFUL::rTCP4-GR* fruits remained similar to that of *ful-1* fruits under uninduced conditions, activation of TCP4 by dexamethasone induction resulted in an increase in fruit length to >10 mm, thus restoring the length to ∼70% of the Col-0 fruit length (**Figure S5e-f**), suggesting a *FUL*-independent role of JAW-TCPs.

*STK* has been shown to promote fruit elongation independent of *FUL*, and its loss of function reduces fruit length to ∼8 mm in the mutant from ∼14 mm in Col-0^41^ (**Figure 5e-f**). To determine the relationship between *JAW-TCP*s and *STK*, we performed genetic interaction studies between the loss-of-function mutants of both genes. Fruits in the *tcp2;3;4;10;stk* plants were significantly shorter than *stk*, whereas the weak allele *ful-6* only marginally reduced the fruit length of *stk* (**Figure 5e-f**). We interpreted all these data as JAW-TCPs promoting fruit length in a *FUL*-independent manner and perhaps in an *STK*-independent manner as well.

## Discussion

Organ growth and morphogenesis usually proceed to completion in continuum, once initiated. In angiosperms, however, some traits such as pistil growth display an intermediate pause and resume only after receiving specific developmental or environmental cues, somewhat akin to the pupal diapause in insects^42^ or embryonic diapause in mammals^43,44^. Upon development and maturation, pistils undergo a growth pause lasting up to a few days, during which the stigmatic papillae remain receptive to the pollen grains. No further pistil growth is observed without fertilization, resulting in an eventual terminal senescence of the unfertilized pistil. However, successful fertilization brings about an end to the pause, triggering rapid growth of the resulting fruit to maturity, perhaps driven by some fertilization-dependent signal from the nascent seeds within the fruit^4^. Whereas the molecular regulation of pistil initiation and patterning have been elucidated in detail, the genes and genetic pathways that trigger post-fertilization fruit growth remain unclear.

Here, we show that the five JAW-TCP transcription factors act upstream to *FUL* to kickstart fruit elongation after fertilization (**Figure 6**). *JAW-TCP*s and *FUL* are expressed in the valve region, where the former bind to the upstream cis-regulatory elements of the *FUL* locus and directly activate its transcription, resulting in the suppression of the valve margin genes, thus allowing the elongation of valve cells throughout the growth process. Both wild-type and mutant fruits (*tcp^Q^*, *jaw-D;ful-6*, *ful-1^Col-0^*) showed typical sigmoidal kinetics, with growth occurring for 10-11 days upon fertilization until maturity at stage 17-B (**Figure 3, S3**). However, the rate of growth was markedly different in the mutants. Though the highest growth was observed at 2-4 DAP in wildtype and in mutant fruits, the rate of growth was much lower for *tcp^Q^* and *jaw-D;ful-6*. Thus, JAW-TCPs affect the rate of fruit growth and not its duration.

**Figure 6.**
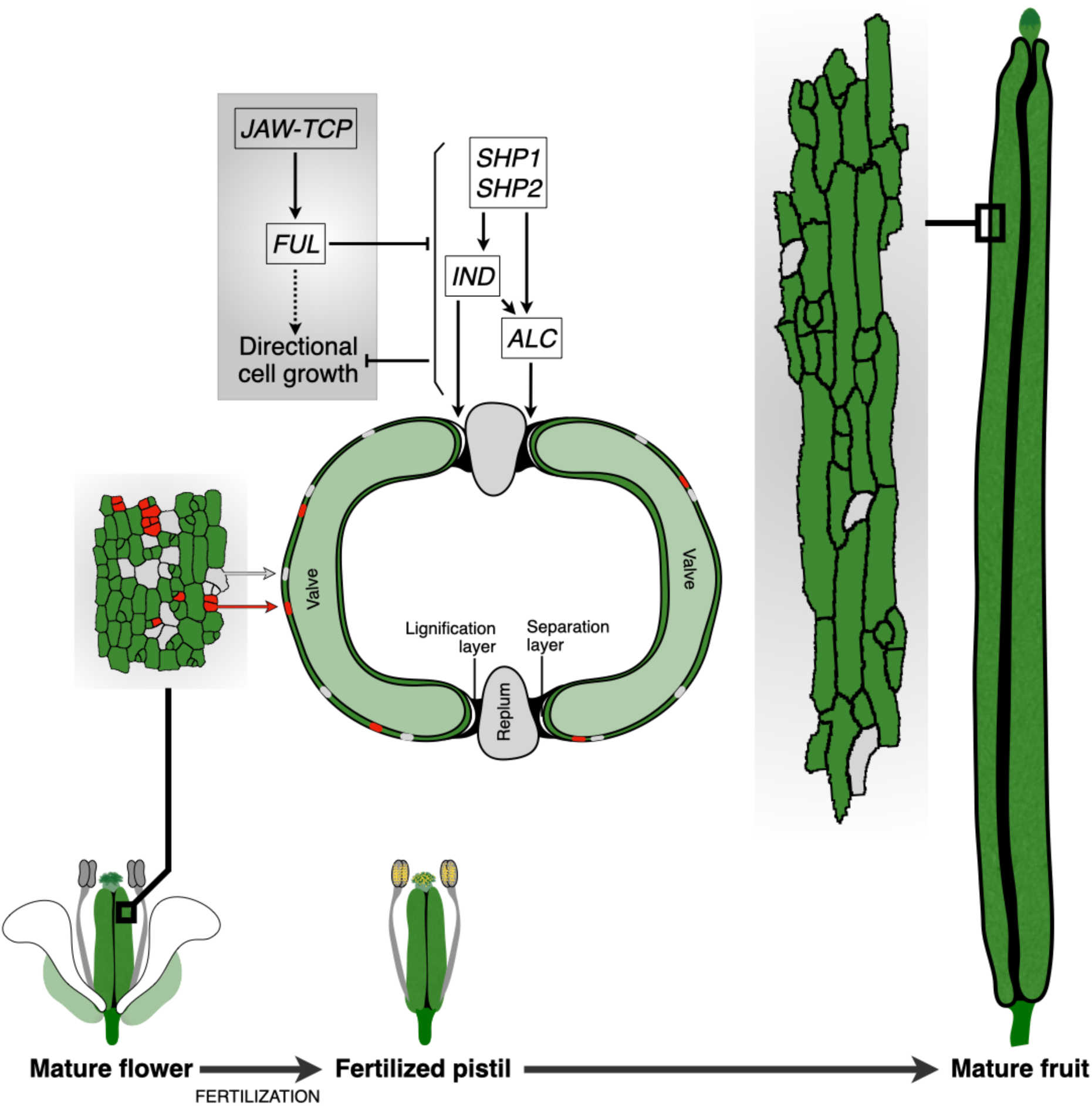
A proposed model for the regulation of fruit elongation by JAW-TCPs. JAW-TCPs directly activate *FUL* transcription (solid arrow), which in turn represses the valve margin genes (*SHP1, SHP2, IND, ALC*), thus restricting their transcripts to the valve margin layers, and allowing anisotropic, vertically oriented growth of the valve cells. In *jaw-tcp* or *ful* loss-of-function mutants, the valve margin genes are ectopically activated in the valves, thereby preventing cell elongation. FUL also promotes valve cell elongation to a smaller extent independent of the valve margin genes (broken arrow). Cartoons of the pavement cells of each orientation type are indicated in green (vertical), grey (intermediate) and red (horizontal).

Fruit elongation in Arabidopsis is driven solely by cell elongation along the length axis, and little change in cell number is observed during this process^4^ (**Figure S4b**). Analysis of the epidermal cell population revealed that the cell anisotropy and its orientation were comparable in the mutant and wild-type fruits soon after pollination (**Figure 4**), suggesting that JAW-TCPs do not regulate pistil growth until fertilization. The proportion of vertically oriented cells in wild-type fruits increased after fertilization until 4 DAP, with a concomitant decrease in the circularity index, triggering fruit elongation. It is worth noting that the maximum increase in the proportion of vertically oriented cells preceded the maximum rate of fruit elongation in the wild type (compare **Figure 4g** with **Figure 3e**), suggesting that the reorientation of cell anisotropy along the fruit growth axis is the first event post-fertilization. This early reorientation of cells was not observed in the mutant fruits (**Figure 4g**), followed by a lack of elongation. This suggests that the earliest function of the *JAW-TCP-FUL* module post-fertilization is to promote directional cell growth along the fruit-length axis.

The JAW-TCPs regulate diverse cellular parameters in plant organs, including cell division, cell expansion, and cell shape^45–47^. There appears to be a difference in their function depending on the type of organs; they suppress cell proliferation in the primordia of lateral organs such as leaves^36,47^ and flowers^30^ whereas their primary function in the primary-axis organs such as hypocotyl^46^ and fruits is to promote directional cell growth (**Figure 1**), even though fruit valves have been considered analogous to leaves^48^. The developmental cue for the direction of cell growth works upstream to *JAW-TCPs*, perhaps mediated by phytohormones such as auxin^8^.

Though *JAW-TCP*s and *FUL* act as a genetic module in promoting fruit elongation, we noted some differences in their mutant phenotypes. Whereas *ful-1*^Col-0^ fruits formed enlarged repla^21^, *tcp^Q^* repla appeared to be normal. Besides, the number of epidermal cells in mature pistil was more in *ful-1*^Col-0^ valves, and their size was relatively smaller compared to Col-0, *tcp^Q^* and *jaw-D;ful-6* (**Figure S4b, d**). These differences arise before fertilization and indicate a *JAW-TCP*- independent role of *FUL* in pistil patterning. Though expressed in the growing pistil^48^, *JAW-TCPs* appear to be involved only in the post-fertilization cell elongation. This is further supported by the lack of any defect in fruit architecture and elongation when JAW-TCPs were downregulated in the valve margin tissue, suggesting that the suppression of margin-specifying genes is *FUL*-dependent.

Although *JAW-TCPs* are expressed during pistil development^29^ where they contribute to the ovule and apical pistil patterning, their effect on *FUL* activation in the ovary wall appears to be inhibited at this stage. Negative regulators of fertilization-independent ovary elongation^49^, such as ARF8^23,24^, may repress JAW-TCP activity in the ovary wall. In the *arf8* mutant, fertilization-independent fruit formation is observed which is dependent on *FUL* as parthenocarpy is abolished in *arf8;ful-1* double mutants^24^. *ARF8* and *JAW-TCPs* have earlier been implicated in the regulation of filament elongation, where the miR319 loss-of-function line phenocopies the loss of mir167-targeted *ARF6* and *ARF8*^50^. It is possible that a converse negative regulation of JAW-TCPs by ARF8 occurs in the ovary wall before fertilization. While direct regulation of *FUL* by ARF8 has not been established, ARF8 may repress JAW-TCP-mediated activation of *FUL* by protein-protein interaction and sequestration, as has been demonstrated earlier for JAW-TCP and CUC proteins in the regulation of leaf serration^51^. The ovular growth signals may remove this suppression following fertilization. Interestingly, *ARF8* and *JAW-TCPs* cell-autonomously function during ovule and seed development, albeit at different stages^23,29,52^. Thus, a possibility exists that the *JAW-TCP-ARF* module initiates and maintains fruit morphogenesis upon successful fertilization.

Fertilization triggers auxin biosynthesis in the ovule, which activates GA metabolism, driving ovary growth^53^; external administration of auxin or gibberellin triggers parthenocarpic fruit formation. JAW-TCPs function upstream of both the hormones in several tissue types; they promote hypocotyl cell elongation by activating the auxin biosynthesis gene *YUC5^46^* in Arabidopsis. In tomato, *LANCEOLATE*, a *JAW-TCP* ortholog, activates GA biosynthesis^54,55^ during leaf development, suppressing leaf complexity. An earlier report showed that, although STK can activate *FUL* transcription in a non-cell autonomous manner^41^, it instead promotes fruit elongation by suppressing cytokinin response, a function also showed by JAW-TCPs in the context of leaf development^35,41^. The synergistic effect of *JAW-TCPs* and *STK* (**Figure 5e-f**) indicates a potential cytokinin-dependent and *FUL*-independent mechanism for fruit elongation post-fertilization.

Fruits in the members of the Brassicaceae family exhibit extensive shape diversity^56^. For instance, while Arabidopsis fruits maintain uniform width along the length forming a cylindrical structure, fruits in *Capsella spp.* exhibit distal widening resulting in a triangular shape^57^. This shape difference across species correlates with altered expression of valve margin genes such as *IND*, which influences cell expansion pattern^57^; abolishing *FUL* function in *Capsella* results in the loss of triangular fruit shape^58^. Early cytoskeleton and cell wall composition changes likely predefine the cell expansion axis, directing fruit morphogenesis. Altered cell anisotropy and its orientation in the *jaw-tcp* fruits make these genes attractive candidates as upstream regulators of fruit shape diversity. Variation in *JAW-TCP* expression patterns may underlie general fruit morphology diversity without affecting tissue morphology, unlike in the mutants of *FUL,* which would also have altered patterning, leading to reduced reproductive success due to the burst of ovary wall and immature seed exposure during growth^21^. Fruit shape variation can be brought about either by the expression diversity of the *JAW-TCP* genes or by changes in the upstream cis-regulatory regions of their downstream target genes, leading to their varied expression.

Our findings in this study highlight the central role of *JAW-TCPs* in regulating directional fruit growth and its shape. The JAW-TCP proteins can be used as a primary model to investigate early events driving fruit wall growth soon after fertilization. It is possible that a signal emanating from the fertilized ovule triggers the JAW-TCP-mediated reorientation of valve cells and initiates fruit elongation. Increased length of the ovary wall cells only adjacent to the fertilized ovules within^4^ suggests that the signal is likely to be mechanical, though the possibility of a chemical signal cannot be excluded. Perhaps a weak *JAW-TCP* mutant such as *tcp2;3;4;10*, which is more poised to show reduced fruit length than the wild type, can be used for a forward mutagenesis screen to identify such a signal if it indeed exists.

## Materials and Methods

### Plant materials

The Columbia-0 (Col-0) ecotype of *Arabidopsis thaliana* was used as the wild-type control. *tcp^Q^* was generated by crossing *tcp2;3;4;10;GR #2*^59^ to *tcp24* (*SALK_077675*), which contained additional T-DNA insertions in *AT5G37160.1* and *AT5G64050.1* that were segregated out in three separate lines F4 plants that were homozygous for *tcp^Q^* (#2, #4 and #14). For all subsequent experiments, *tcp^Q^* #4 was used. *ful-1* was obtained from ABRC (CS3759) and backcrossed to Col-0 four times before being used for all experiments as *ful-1*^Col-0^. *jaw-D;GR* #1^46^ and *tcp2;3;4;10;GR* #2 were crossed to *ful-6* to obtain *tcp2;3;4;10;ful-6* and *jaw-D;ful-6*. The plasma membrane reporter containing *tcp2;3;4;10;24*, *jaw-D;ful-6* and *ful-1*^Col-0^ were generated by crossing each mutant to *pUBQ10::ACYL-YFP*^34^. All other higher-order mutant combinations were generated by crossing the respective homozygous parents and establishing the homozygous lines in the F2 and later generations.

### Generation of transgenic lines

A 2.2 kb DNA fragment corresponding to *proFUL* and a 2.15 kb DNA fragment corresponding to *proSHP2* were cloned into *pDONR-P4P1R*. A subsequent LR reaction (11791020, Invitrogen, USA) was carried out with *pDONR-P4P1R-proFUL* or *proSHP2* and the destination *R4L1pGWB532* vector to generate *proFUL::G3GFP-GUS* (referred to as *proFUL::GUS*) and *proSHP2::G3GFP-GUS* (referred to as *proSHP2::GUS*). The *FUL* cDNA was cloned into *pDONR221* and combined with *pDONR-P4P1R-proTCP4* into *R4pGWB507* to obtain *pTCP4::FUL*. *pDONR-P4P1R-proFUL* and *pDONR221-rTCP4-GR*^59^ were combined into *R4pGWB507* to obtain *proFUL::rTCP4-GR*. *pDONR-P4P1R-proSHP2* and *pDONR221-MIR319A*^59^ were combined into *R4pGWB507* to generate *proSHP2::MIR319A*. All cloning primers are listed in Supplementary Table 1.

Chemically competent *E. coli* DH5α cells were transformed with the BP and LR clonase reaction products and plated on respective antibiotic selection plates. Liquid cultures were prepared from colonies which grew on the selection plates. Plasmids were purified from the liquid cultures QIAprep Spin Miniprep Kit (Qiagen, Germany). Plasmids were validated by restriction digestion followed by Sanger sequencing. Electrocompetent *Agrobacterium tumifaciens* GV3101 cells were transformed with the binary constructs, and positive colonies were confirmed by colony PCR. Constructs were then integrated into the Col-0 genome by the *Agrobacterium tumefaciens*-mediated floral dip method^60^ using 50 μl/ litre Silwet L-77. Seeds were harvested from primary transformants, and the T1 transformants were selected using 20 mg/l hygromycin-supplemented MS plates.

### Plant growth conditions and chemical treatments

Seeds were treated with a solution of 70% ethanol and 0.05% SDS to surface sterilize them and kept on a rocker for 10-15 minutes, following which they were washed 2-3 times with 100% ethanol, dried on Whatman filter papers and sown on 1x MS plates. Sown seeds were stratified for three days in the dark at 4°C and then shifted to a plant growth chamber (GC-1000 TLH, JEIO TECH, KOREA). The growth chamber was set to 22°C with a light intensity of 120 µmol/m^2^ for a long day photoperiod of 16 hours followed by 8 hours of darkness. Dexamethasone treatment was done using 12 µM dexamethasone solution or mock ethanol solvent and sprayed onto inflorescences.

### β-glucuronidase (GUS) assay and imaging

Gynoecia were harvested and fixed in 90% ice-cold acetone for 15-20 min at room temperature, rinsed with GUS buffer (0.5 M Sodium phosphate buffer, pH 7.2; 10% Triton X-100; 100 mM potassium ferrocyanide; 100 mM potassium ferricyanide) for two minutes, placed in fresh buffer containing 1 mM X-Gluc (Thermo Scientific, USA), and incubated at 37°C for 12-16 hours, before clearing with 70% ethanol at 12-16 hour intervals until the samples were clear. Samples were then mounted in water, and pictures were taken using a Leica (Germany) EZ4E stereo zoom microscope.

### RNA isolation and RT-qPCR

40-50 mg of gynoecia corresponding to stages 14 and 15 were collected for RNA isolation using TRIzol (Sigma-Aldrich, USA). 10 µg of the total RNA isolated was treated with DNase enzyme (1 Unit/µL-Thermo Scientific, USA), and 1.5 µg of the product was converted to cDNA using the RevertAid RT reverse transcription kit (K1691, Thermo Scientific, USA). 25 ng of cDNA was used as a template for quantitative PCR using DyNAmo ColorFlash SYBR Green qPCR Kit (F416L, Thermo Scientific, USA) in the QuantStudio Real-Time PCR System (Quant5 or Quant 6, Applied Biosystems, USA). Raw CT values were obtained from the built-in Applied Biosystems software. ΔΔCT values were determined after normalization with internal control. Fold change was calculated as previously reported^61^. All RT-qPCR primers are listed in Supplementary Table 1.

### Electrophoretic Mobility Shift Assay (EMSA)

End-labelled oligonucleotides (Supplementary Table 1) corresponding to the class-II TCP consensus binding site and +/- 12 bp flanking it were prepared with [γ-32P]-ATP using T4 polynucleotide kinase (Thermo Scientific, USA). A 15 µL binding reaction containing an oligonucleotide probe, 1X binding buffer, and approximately 2 mg of crude recombinant protein bacterial lysate was prepared as previously described^46^ and incubated at room temperature for 30-40 min followed by resolving on a 9% native polyacrylamide gel in 0.5X TBE buffer. A monoclonal α-HIS antibody (Sigma-Aldrich; H1029) was used at 30X final dilution in the binding buffer for the super-shift assay. A 1x or 10x concentration of unlabelled probe was used in a competition assay to compete with the labelled probe. The gels were autoradiographed using a Phosphor imager (GE Typhoon FLA 9500, GE).

### Chromatin immunoprecipitation (ChIP) assay

ChIP was performed as mentioned earlier^38^ with a few modifications. Gynoecia comprising stages 14-15 (75-100 mg) were harvested per sample. Isolated chromatin was sonicated using the Bioruptor Plus (Diagenode, USA) system set at low mode and 45 cycles of 30 sec ON and 45 sec OFF to obtain sheared chromatin from 500 bp to 1kb. 2% sheared chromatin was set aside as input for later control experiments. 3 μL anti-FLAG antibody (F1804, Sigma-Aldrich, USA) precipitated the cross-linked complexes (4 hours at 4°C on a rotary shaker). Immunoprecipitated DNA was purified with the QIAquick PCR purification kit (28104, Qiagen, Germany). The purified DNA was resuspended in 100 μL of elution buffer, and qPCR was set up with 3μL DNA as a template per reaction. Fold enrichment of TCP4 in α-FLAG samples was expressed relative to % input. All ChIP-qPCR primers are listed in Supplementary table 1.

### Yeast two-hybrid (Y2H) assay

The prey constructs were selected from an existing assorted library where 1956 transcription factors are fused with the GAL4 activation domain and cloned in the pDEST22 vector^62^. The reporter strain AH109, bait vector pDEST32 (with LEU marker) and the prey vector pDEST22 (with TRP marker) were used for the assay. AH109 yeast cells were transformed with the LiOAc/ssDNA/PEG method (Yeast Protocol Handbook, Clonetech, CA) and plated on synthetic dropout (SD) media lacking Trp and Leu to identify the co-transformants. A single colony generated a primary culture in a liquid SD media lacking Trp and Leu. The OD from each bait-prey culture was diluted to 0.3 to serve as undiluted cultures for the Y2H assay. From the undiluted culture, dilutions up to 10^-4^ were prepared and then spotted on SD media lacking Trp, Leu, Ade and His Y2H assay plates were incubated at 30°C in an incubator and checked for growth after 2–3 days.

### Generation of CRISPR-edited lines

Guide RNAs (gRNA) targeting the BS2 and B3-4 TCP4-binding sites on the *FUL* locus were designed using CRISPOR^63^. As previously reported^64,65^, BS2 and BS3-4 gRNA oligos were cloned into pAGM55273 by a BsaI-Golden Gate reaction (BsaI-HF®v2, #R3733S, New England Biolabs, USA). Similarly, gRNA targeting *SHP1*, *SHP2*, *IND* and *ALC* were designed and cloned into shuttle vectors M1, M2, M3 and M4E followed by assembly into pDGE347. All CRISPR-related oligos are listed in Supplementary table 1. Chemically competent *E. coli* DH5α cells were transformed with the reaction products and plated on respective antibiotic selection plates, 2-4 colonies were tested by restriction digestion and validated by Sanger sequencing. *Agrobacterium tumifaciens* GV3101 cells were transformed with binary constructs by electroporation, and a single colony was confirmed by colony PCR. Constructs were then integrated into the respective lines by the *Agrobacterium tumefaciens*-mediated floral dip method^60^. T1 seeds were collected and dried. Positive transformants were selected by screening for RFP signals in the seed coat in an Olympus stereo zoom microscope fitted with an RFP fluorophore set. T1 plants were screened for phenotypic defects in fruit elongation and tested for mutations in the putative JAW-TCP binding sites in *FUL* or the *SHP1*/*SHP2*/*IND*/*ALC* genes by Sanger sequencing. A biallelic/heterozygous mutation was confirmed in the T1 generation, and a homozygous Cas9-free T2 mutant was obtained and confirmed by Sanger sequencing.

### Confocal microscopy

Pistils and fruits were collected at indicated time points or stages of development and mounted in water on a glass slide, and the coverslip was sealed with clear nail polish. Imaging was done in the Olympus Fluoview FV10i system (Japan) in the LSM mode. YFP signals were excited at 473 nm, and emissions were collected at 527 nm.

### Microtome sectioning

Tissues were first collected into scintillation vials, fixed in FAA (absolute alcohol-50%, acetic acid-5%, formaldehyde-10%, water-35%) overnight, then passed through an ethanol series of 50%, 60%, 70%, 80%, 90% (all prepared in water), and absolute ethanol. Tissues were then passed through a xylene series of 25%, 50%, 75% (all prepared in absolute ethanol) and 100% xylene. 100 chips of Paraplast Plus were added to the solution and left overnight. The xylene-Paraplast Plus solution was transferred to a dry bath at 65°C the following day. The solution was replaced with a molten Paraplast Plus solution 4 times in intervals of at least 4 hours. The tissue-containing solution was then poured into a mould and allowed to solidify. Thin sections were obtained at 8 µm thickness and mounted onto water-containing glass slides on a slide warmer set at 42°C. After drying, the slides were reversed through the xylene and ethanol series, stained with 1% Toluidene blue or 2% phloroglucinol, and returned to 100% xylene before mounting in the DPX mountant.

### Statistical analysis

All graphical representations and statistical analyses were performed in R^66–68^. In plots where statistical analysis has been included, similar groups are indicated by similar alphabets, while the difference in letters indicates statistical significance.

## Accession numbers

TAIR (The Arabidopsis Information Resource, www.arabidopsis.org) accession numbers of the significant genes used in this study are *AT4G18390* (*TCP2*), *AT1G53230* (*TCP3*), *AT3G15030* (*TCP4*), *AT2G31070* (*TCP10*), *AT4G23713* (*MIR319A*), *AT1G30210* (*TCP24*), *AT2G40670* (*ARR16*), *AT5G60910* (*FUL*), *AT3G58780* (*SHP1*), *AT2G42830* (*SHP2*), *AT5G67110* (*ALC*), *AT4G00120* (*IND*) and *AT4G09960* (*STK*).

## Competing Interests

The authors declare no conflict of interest.

## Funding

This work was supported by fellowships from the Ministry of Education (A.N.S.), Department of Biotechnology (S.B.), Council of Scientific and Industrial Research (K.C., M.K.), and Anusandhan National Research Foundation (JC Bose Fellowship to U.N.) from Government of India. Authors thank the Department of Science & Technology for Improvement of S&T Infrastructure (DST-FIST, No. SR/FST/LSII-044/2016 dated 15.12.2016) and the Department of Biotechnology (DBT)-IISc Partnership Program Phase-II at IISc (sanction No. BT/PR27952/INF/22/212/ 2018) for the funding and infrastructure support. The funders had no role in study design, data collection and analysis, decision to publish, or preparation of the manuscript.

## Author contributions

A.N.S. initiated the project, designed & performed most experiments, analysed & interpreted the results, organized the figures, wrote the first draft of the manuscript and contributed to its finalization. K. and A.V. participated in collecting data from most experiments and carried out the Crispr/Cas9 editing of BS3 in *FUL* (mostly A.V., with inputs from A.N.S) and valve margin-specifying genes (K.). A.V. and N.S.S.S. collected data from the confocal experiments with the help of S.B. and K.C. A.J.P. cloned *pTCP4::FUL* and *pFUL::rTCP4-GR*. M.K. and D.P. were involved in the initial experiments and establishing *jaw-D;ful-6,* and *tcp^Q^*. S.B. carried out all the RT-qPCR, ChIP-qPCR and EMSA experiments and contributed to the finalization of the manuscript. U.N. contributed to designing experiments and data interpretation, guided the authors, corrected and finalised the manuscript.

## Acknowledgements

Naveen Shankar (IISc Bangalore, India) is acknowledged for generating *pDONR-P4P1R-pTCP4* and Kavita Babu (IISc Bangalore, India) is acknowledged for helping with the Olympus stereo zoom fluorescent stereomicroscope. We thank Sachin Kotak, Yadukrishnan Premachandran and Naveen Shankar of IISc Bangalore, India, for helpful discussions during the preparation of this manuscript. Thanks to Juan Ignacio Ezquer Garin (UNIMI, Italy) for the *stk* mutant seeds, Daniel Kierzkowski (Université de Montréal) for *pUBQ10::ACYL-YFP* seeds, Martin Yanofsky (UC San Diego) for the *ful-6* mutant seeds and Tsuyoshi Nakagawa (Shimane University) for the *R4pGWB507* and *R4L1pGWB532* vectors.

## Notes

### Competing Interest Statement

The authors have declared no competing interest.

## References

1. Wiens, D. et al. Reproductive success, spontaneous embryo abortion, and genetic load in flowering plants. Oecologia 71, 501–509 (1987).

2. Simpson, M. G. Plant Systematics, Third Edition. Plant Systematics, Third Edition 1–761 (2019) doi:10.1016/C2015-0-04664-0.

3. Smyth, D. R., Bowman, J. L. & Meyerowitz, E. M. Early flower development in Arabidopsis. Plant Cell 2, 755–767 (1990).

4. Ripoll, J. J. et al. Growth dynamics of the Arabidopsis fruit is mediated by cell expansion. Proc Natl Acad Sci U S A 116, 25333–25342 (2019).

5. Arnaud, N. et al. Gibberellins Control Fruit Patterning in Arabidopsis Thaliana. Genes Dev 24, 2127–2132 (2010).

6. Feys, BJF., Benedetti, C. E., Penfold, C. N. & Turner, J. G. Arabidopsis Mutants Selected for Resistance to the Phytotoxin Coronatine Are Male Sterile, Insensitive to Methyl Jasmonate, and Resistant to a Bacterial Pathogen. Plant Cell 6, 751 (1994).

7. Marsch-Martínez, N. et al. The Role of Cytokinin during Arabidopsis Gynoecia and Fruit Morphogenesis and Patterning. The Plant Journal 72, 222–234 (2012).

8. Nemhauser, J. L., Feldman, L. J. & Zambryski, P. C. Auxin and ETTIN in Arabidopsis gynoecium morphogenesis. Development 127, 3877–3888 (2000).

9. Rong, D., Luo, N., Mollet, J. C., Liu, X. & Yang, Z. Salicylic Acid Regulates Pollen Tip Growth through an NPR3/NPR4-Independent Pathway. Mol Plant 9, 1478–1491 (2016).

10. Ye, Q. et al. Brassinosteroids control male fertility by regulating the expression of key genes involved in Arabidopsis anther and pollen development. Proc Natl Acad Sci U S A 107, 6100–6105 (2010).

11. Bowman, J. L., Smyth, D. R. & Meyerowitz, E. M. Genes directing flower development in Arabidopsis. Plant Cell 1, 37 (1989).

12. Colombo, M. et al. A new role for the SHATTERPROOF genes during Arabidopsis gynoecium development. Dev Biol 337, 294–302 (2010).

13. Pinyopich, A. et al. Assessing the redundancy of MADS-box genes during carpel and ovule development. Nature 2003 424:6944 424, 85–88 (2003).

14. Ferrándiz, C., Liljegren, S. J. & Yanofsky, M. F. Negative regulation of the SHATTERPROOF genes by FRUITFULL during Arabidopsis fruit development. Science 289, 436–438 (2000).

15. Roeder, A. H. K., Ferrándiz, C. & Yanofsky, M. F. The role of the REPLUMLESS homeodomain protein in patterning the Arabidopsis fruit. Current Biology 13, 1630–1635 (2003).

16. Ferrándiz, C., Gu, Q., Martienssen, R. & Yanofsky, M. F. Redundant regulation of meristem identity and plant architecture by FRUITFULL, APETALA1 and CAULIFLOWER. Development 127, 725–734 (2000).

17. Liljegren, S. J. et al. SHATTERPROOF MADS-box genes control seed dispersal in Arabidopsis. Nature 404, 766–770 (2000).

18. Roeder, A. H. K. & Yanofsky, M. F. Fruit Development in Arabidopsis. Arabidopsis Book 2006, (2006).

19. Liljegren, S. J. et al. Control of fruit patterning in Arabidopsis by INDEHISCENT. Cell 116, 843–853 (2004).

20. Rajani, S. & Sundaresan, V. The Arabidopsis myc/bHLH gene ALCATRAZ enables cell separation in fruit dehiscence. Current Biology 11, 1914–1922 (2001).

21. Gu, Q., Ferrándiz, C., Yanofsky, M. F. & Martienssen, R. The FRUITFULL MADS-box gene mediates cell differentiation during Arabidopsis fruit development. Development 125, 1509–1517 (1998).

22. Vries, F. D., Laar, H. H. & Chardon, M. Bioenergetics of growth of seeds, fruits and storage organs. (1983).

23. Goetz, M., Vivian-Smith, A., Johnson, S. D. & Koltunow, A. M. AUXIN RESPONSE FACTOR8 Is a Negative Regulator of Fruit Initiation in Arabidopsis. Plant Cell 18, 1873–1886 (2006).

24. Vivian-Smith, A., Luo, M., Chaudhury, A. & Koltunow, A. Fruit Development Is Actively Restricted in the Absence of Fertilization in Arabidopsis. Development 128, 2321–2331 (2001).

25. Vivian-Smith, A. & Koltunow, A. M. Genetic Analysis of Growth-Regulator-Induced Parthenocarpy in Arabidopsis. Plant Physiol 121, 437 (1999).

26. Dinneny, J. R., Weigel, D. & Yanofsky, M. F. A Genetic Framework for Fruit Patterning in Arabidopsis Thaliana. Development 132, 4687–4696 (2005).

27. Goethe, J. W. von. Versuch Die Metamorphose Der Pflanzen Zu Erklären. Versuch die Metamorphose der Pflanzen zu erklären (bey Carl Wilhelm Ettinger, Gotha, 1790). doi:10.5962/bhl.title.127448.

28. Palatnik, J. F. et al. Control of leaf morphogenesis by microRNAs. Nature 2003 425:6955 425, 257–263 (2003).

29. Lan, J. et al. Arabidopsis TCP4 transcription factor inhibits high temperature-induced homeotic conversion of ovules. Nature Communications 2023 14:1 14, 1–19 (2023).

30. Nag, A., King, S. & Jack, T. miR319a Targeting of TCP4 Is Critical for Petal Growth and Development in Arabidopsis. Proceedings of the National Academy of Sciences 106, 22534–22539 (2009).

31. Wang, Y. et al. Arabidopsis transcription factor TCP4 controls the identity of the apical gynoecium. Plant Cell 36, 2668–2688 (2024).

32. Zheng, X., et al. Arabidopsis transcription factor TCP4 represses chlorophyll biosynthesis to prevent petal greening. Plant Commun 3, (2022).

33. Bowman, J. L.. Arabidopsis: an atlas of morphology and development. 450 (1994).

34. Willis, L. et al. Cell size and growth regulation in the Arabidopsis thaliana apical stem cell niche. Proc Natl Acad Sci U S A 113, E8238–E8246 (2016).

35. Efroni, I. et al. Regulation of Leaf Maturation by Chromatin-Mediated Modulation of Cytokinin Responses. Dev Cell 24, 438–445 (2013).

36. Challa, K. R., Rath, M. & Nath, U. The CIN-TCP Transcription Factors Promote Commitment to Differentiation in Arabidopsis Leaf Pavement Cells via Both Auxin-Dependent and Independent Pathways. PLoS Genet 15, e1007988 (2019).

37. Kosugi, S. & Ohashi, Y. DNA Binding and Dimerization Specificity and Potential Targets for the TCP Protein Family. The Plant Journal 30, 337–348 (2002).

38. Kubota, A. et al. TCP4-Dependent Induction of CONSTANS Transcription Requires GIGANTEA in Photoperiodic Flowering in Arabidopsis. PLoS Genet 13, e1006856 (2017).

39. Alvarez-Buylla, E. R. et al. Flower Development. 10.1199/tab.0127 2010, e0127 (2010).

40. Roeder, A. H. K. & Yanofsky, M. F. Fruit Development in Arabidopsis. The Arabidopsis Book / American Society of Plant Biologists 4, e0075 (2006).

41. Di Marzo, M. et al. SEEDSTICK Controls Arabidopsis Fruit Size by Regulating Cytokinin Levels and FRUITFULL. Cell Rep 30, 2846–2857.e3 (2020).

42. J A Powell. Longest Insect Dormancy: Yucca Moth Larvae (Lepidoptera: Prodoxidae) Metamorphose After 20, 25, and 30 Years in Diapause. Ann Entomol Soc Am 94, 677–680 (2001).

43. Iyer, D. P. et al. mTOR activity paces human blastocyst stage developmental progression. Cell 187, 6566–6583.e22 (2024).

44. Ptak, G. E. et al. Embryonic Diapause Is Conserved across Mammals. PLoS One 7, e33027 (2012).

45. Vadde, B. V. L., Challa, K. R. & Nath, U. The TCP4 Transcription Factor Regulates Trichome Cell Differentiation by Directly Activating GLABROUS INFLORESCENCE STEMS in Arabidopsis Thaliana. The Plant Journal 93, 259–269 (2018).

46. Challa, K. R., Aggarwal, P. & Nath, U. Activation of YUCCA5 by the Transcription Factor TCP4 Integrates Developmental and Environmental Signals to Promote Hypocotyl Elongation in Arabidopsis. Plant Cell 28, 2117–2130 (2016).

47. Rodriguez, R. E. et al. Control of Cell Proliferation in Arabidopsis Thaliana by microRNA miR396. Development 137, 103–112 (2010).

48. Girin, T., Sorefan, K. & Østergaard, L. Meristematic sculpting in fruit development. J Exp Bot 60, 1493–1502 (2009).

49. Fuentes, S. & Vivian-Smith, A. Fertilization and Fruit Initiation. Annual Plant Reviews online 107–171 (2018) doi:10.1002/9781119312994.APR0411.

50. Rubio-Somoza, I. & Weigel, D. Coordination of Flower Maturation by a Regulatory Circuit of Three MicroRNAs. PLoS Genet 9, 1003374 (2013).

51. Rubio-Somoza, I. et al. Temporal Control of Leaf Complexity by miRNA-Regulated Licensing of Protein Complexes. Current Biology 24, 2714–2719 (2014).

52. Kong, Q. et al. The Function of the WRI1-TCP4 Regulatory Module in Lipid Biosynthesis. Plant Signal Behav 15, 1812878 (2020).

53. Nitsch, J. P. Plant Hormones in the Development of Fruits. 10.1086/398643 27, 33–57 (1952).

54. Ori, N. et al. Regulation of LANCEOLATE by miR319 is required for compound-leaf development in tomato. Nature Genetics 2007 39:6 39, 787–791 (2007).

55. Yanai, O., Shani, E., Russ, D. & Ori, N. Gibberellin partly mediates LANCEOLATE activity in tomato. Plant J 68, 571–582 (2011).

56. Łangowski, Ł., Stacey, N. & Østergaard, L. Diversification of fruit shape in the Brassicaceae family. Plant Reproduction 2016 29:1 29, 149–163 (2016).

57. Dong, Y. et al. Regulatory Diversification of INDEHISCENT in the Capsella Genus Directs Variation in Fruit Morphology. Curr Biol 29, 1038–1046.e4 (2019).

58. Eldridge, T. et al. Fruit shape diversity in the Brassicaceae is generated by varying patterns of anisotropy. Development 143, 3394–3406 (2016).

59. Shankar, N., Sunkara, P. & Nath, U. A double-negative feedback loop between miR319c and JAW-TCPs establishes growth pattern in incipient leaf primordia in Arabidopsis thaliana. PLoS Genet 19, e1010978 (2023).

60. Clough, S. J. & Bent, A. F. Floral dip: a simplified method for Agrobacterium-mediated transformation of Arabidopsis thaliana. Plant J 16, 735–743 (1998).

61. Livak, K. J. & Schmittgen, T. D. Analysis of relative gene expression data using real-time quantitative PCR and the 2-ΔΔCT method. Methods 25, 402–408 (2001).

62. Pruneda-Paz, J. L. et al. A Genome-Scale Resource for the Functional Characterization of Arabidopsis Transcription Factors. Cell Rep 8, 622–632 (2014).

63. Concordet, J. P. & Haeussler, M. CRISPOR: intuitive guide selection for CRISPR/Cas9 genome editing experiments and screens. Nucleic Acids Res 46, W242–W245 (2018).

64. Grützner, R. et al. High-efficiency genome editing in plants mediated by a Cas9 gene containing multiple introns. Plant Commun 2, 100135 (2021).

65. Stuttmann, J. et al. Highly efficient multiplex editing: one-shot generation of 8× Nicotiana benthamiana and 12× Arabidopsis mutants. Plant J 106, 8–22 (2021).

66. R Core Team & R Foundation for Statistical Computing. R: A Language and Environment for Statistical Computing. https://www.R-project.org/ Preprint at (2024).

67. Wickham, H. et al. Welcome to the Tidyverse. J Open Source Softw 4, 1686 (2019).

68. Hadley Wickham. Ggplot2: Elegant Graphics for Data Analysis. (Springer-Verlag New York, 2016).

