## Supplementary figures and table for "The *JAW-TCP-FUL* genetic axis triggers an early reorientation of cell anisotropy to initiate and drive fertilization-dependent fruit elongation in Arabidopsis"

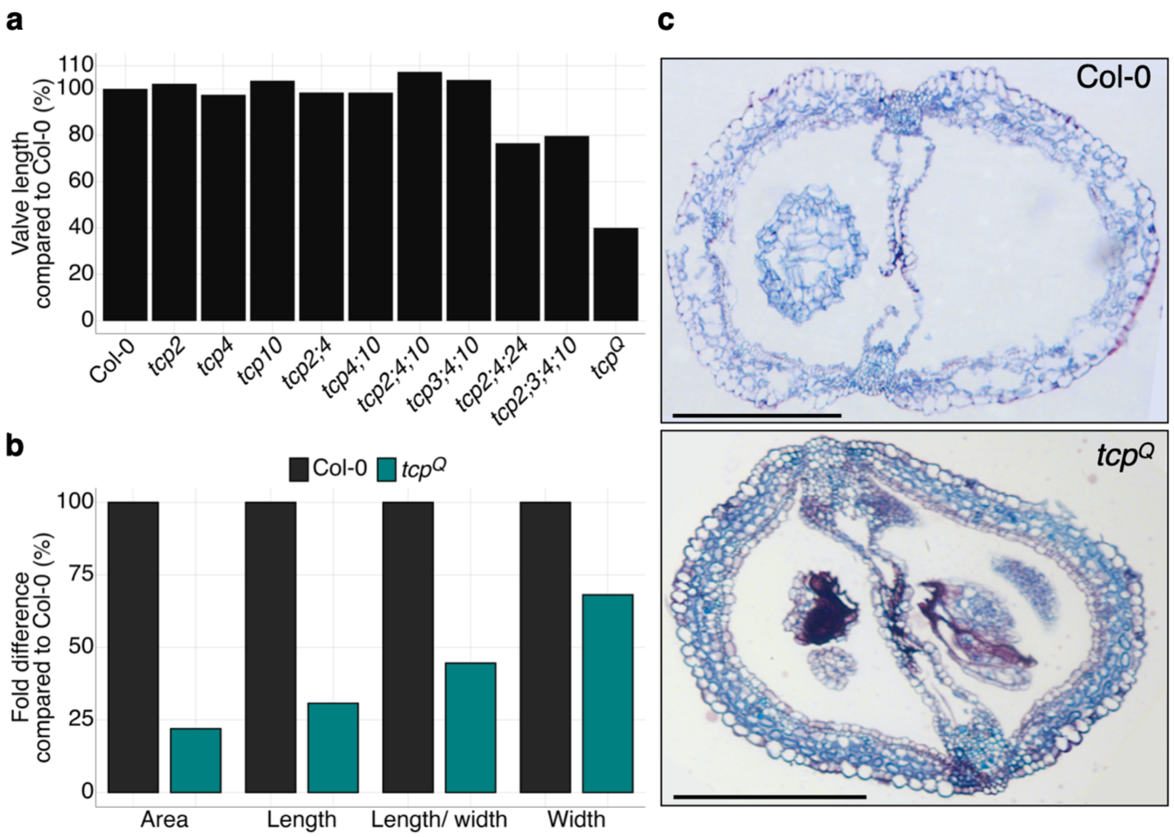


**Figure S1. Relative change in the morphometric parameters and histology of *jaw-tcp* mutants.**

(a, b) Fold difference (%) of the average valve length of fruits (a) and valve epidermal cell parameters (b) relative to Col-0. (c) Toluidine blue-stained transverse sections of mature Col-0 and *tcp*$^{Q}$ fruits. Scale bar, 200 µm (c).

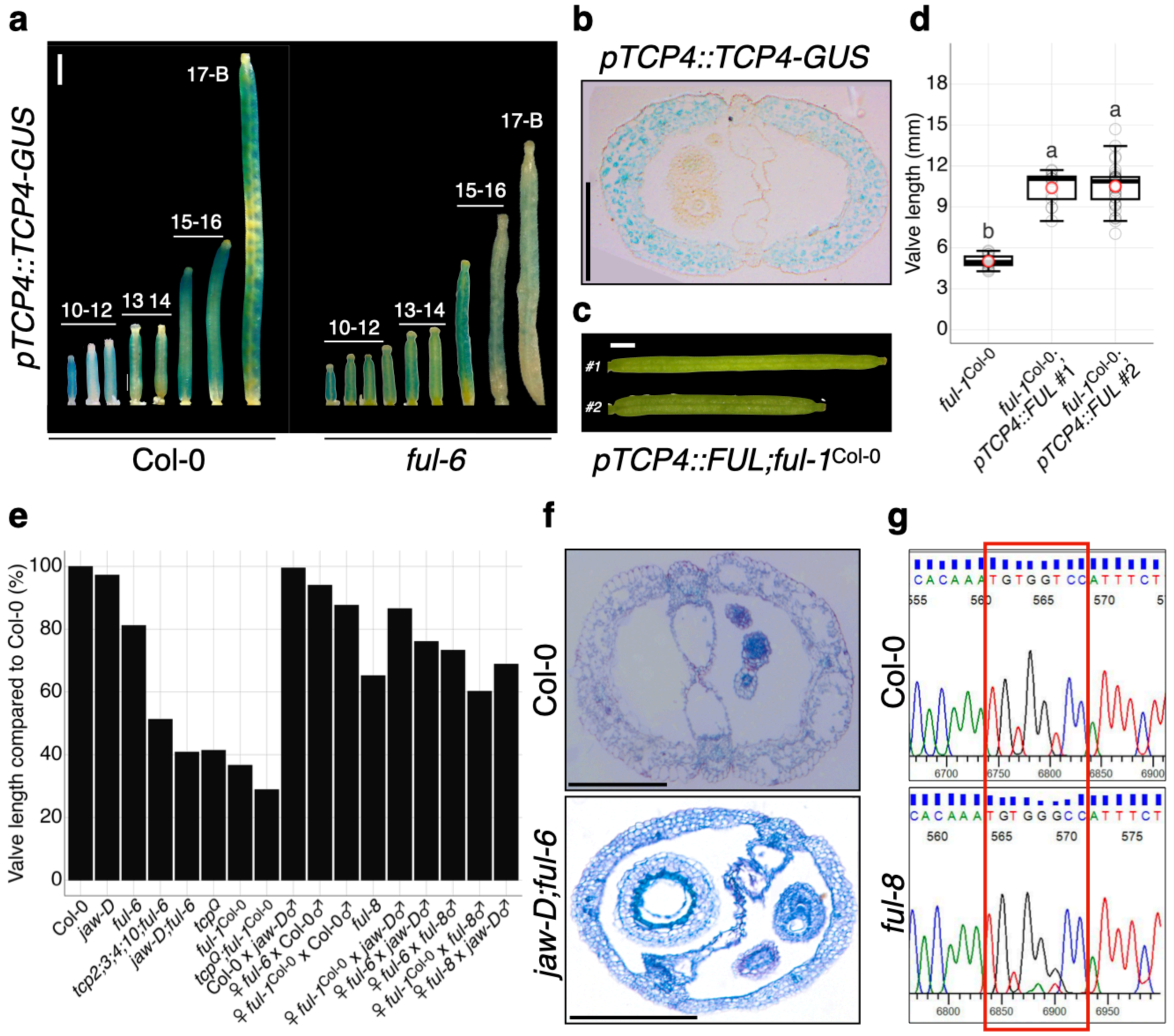


**Figure S2.** ***FUL*** **function is impaired in mutants of** ***JAW-TCPs*** **genes.**

(a) *pTCP4::TCP4-GUS* reporter analysis of Col-0 and *ful-6* gynoecia. Numbers above the fruits indicate stages of growth[3]. (b) Image of a transverse section through a stage 15 *pTCP4::TCP4-GUS* fruit in the Col-0 background displaying GUS staining in the valves[31]. (c, d) Images of mature fruits (c) from two independent T1 *ful-1;pTCP4::FUL* transgenic lines (#1 and #2) and the valve lengths (d) of 13-32 fruits of the indicated genotypes collected from at least three plants per grown batch across three independent batches. Open grey and open red circles indicate individual fruit valve length and average valve length, respectively. The difference in letters marked above the boxplots in (d) between genotypes indicates the statistical significance of $p<0.05$ as measured by a one-way ANOVA followed by Tukey's HSD. (e) Fold difference (%) of the average valve length of fruits indicates genotypes relative to Col-0. (f) Toluidine blue-stained transverse sections of stage 17-A Col-0 and *jaw-D;ful-6* fruits. (g) Sanger sequencing results of Col-0 and *ful-8* upstream regulatory sequences with the predicted JAW-TCP binding site (BS3) boxed in red. The T→G transversion in *ful-8* is boldfaced in the sequences below the boxes. Each boxplot in (d) indicates the second and third quartile of data points, with the horizontal bar representing the median. The upper and the lower whiskers indicate either the highest/ lowest value or ±1.5 times the interquartile range. Scale bar, 1 mm (a, c), 200 µm (b, f).

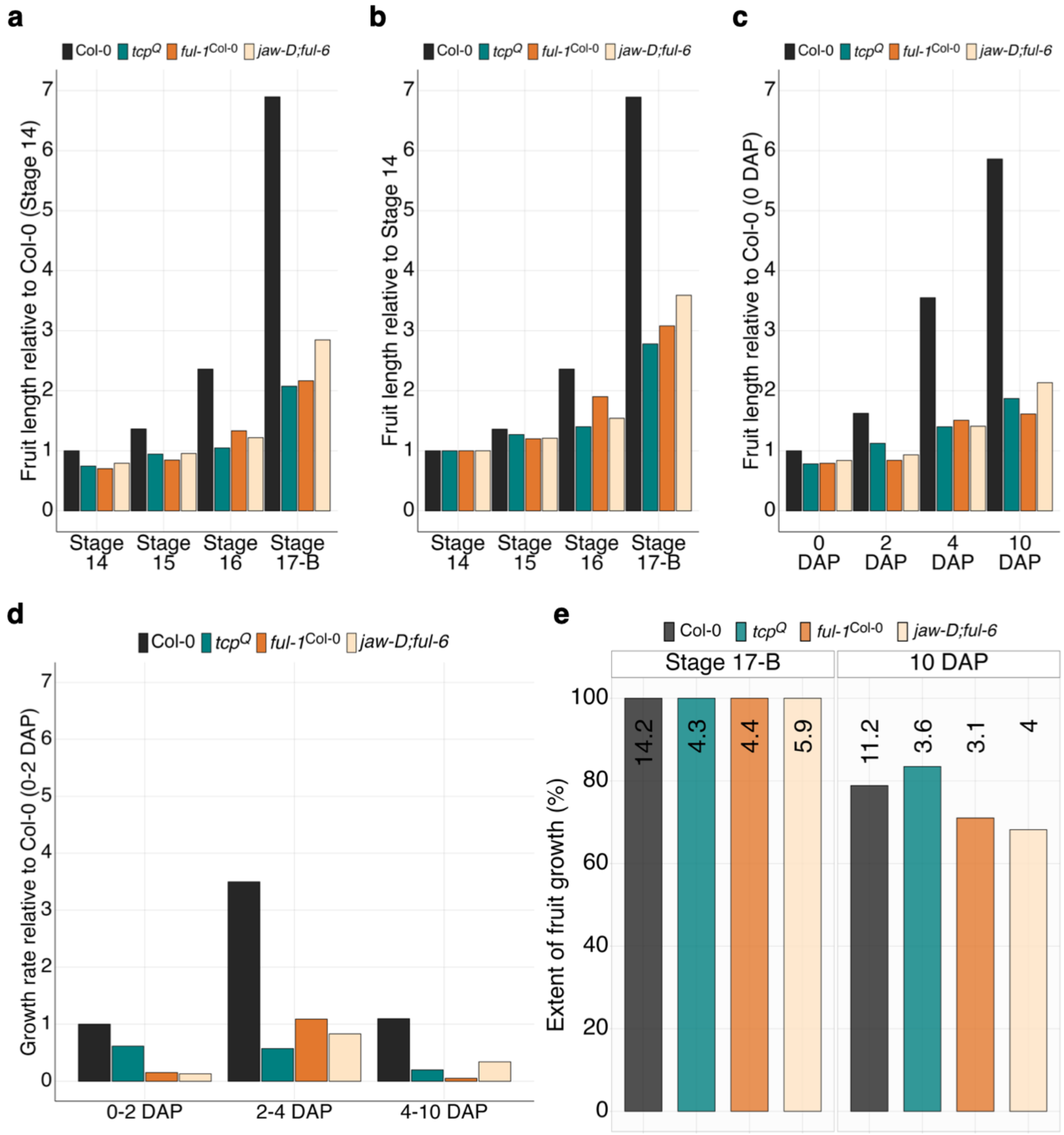


**Figure S3. The rate of fruit elongation and not its duration is impaired in the absence of *JAW-TCPs* and *FUL* function.**

(a-c) Fold difference of the average valve length of fruits of indicated genotypes compared to stage 14 Col-0 (a), compared to the respective genotype at growth stage 14 (b), and compared to Col-0 at 0 days after pollination (DAP) (c). (d) The relative fruit growth rate between 0-10 DAP in each genotype compared to the Col-0 value between 0-2 DAP. (e) Percent of fruit length attained at 10 DAP in indicated genotypes relative to their respective mature lengths at stage 17-B. Numbers at the top of each bar indicate absolute length at the respective stage/DAP in mm.

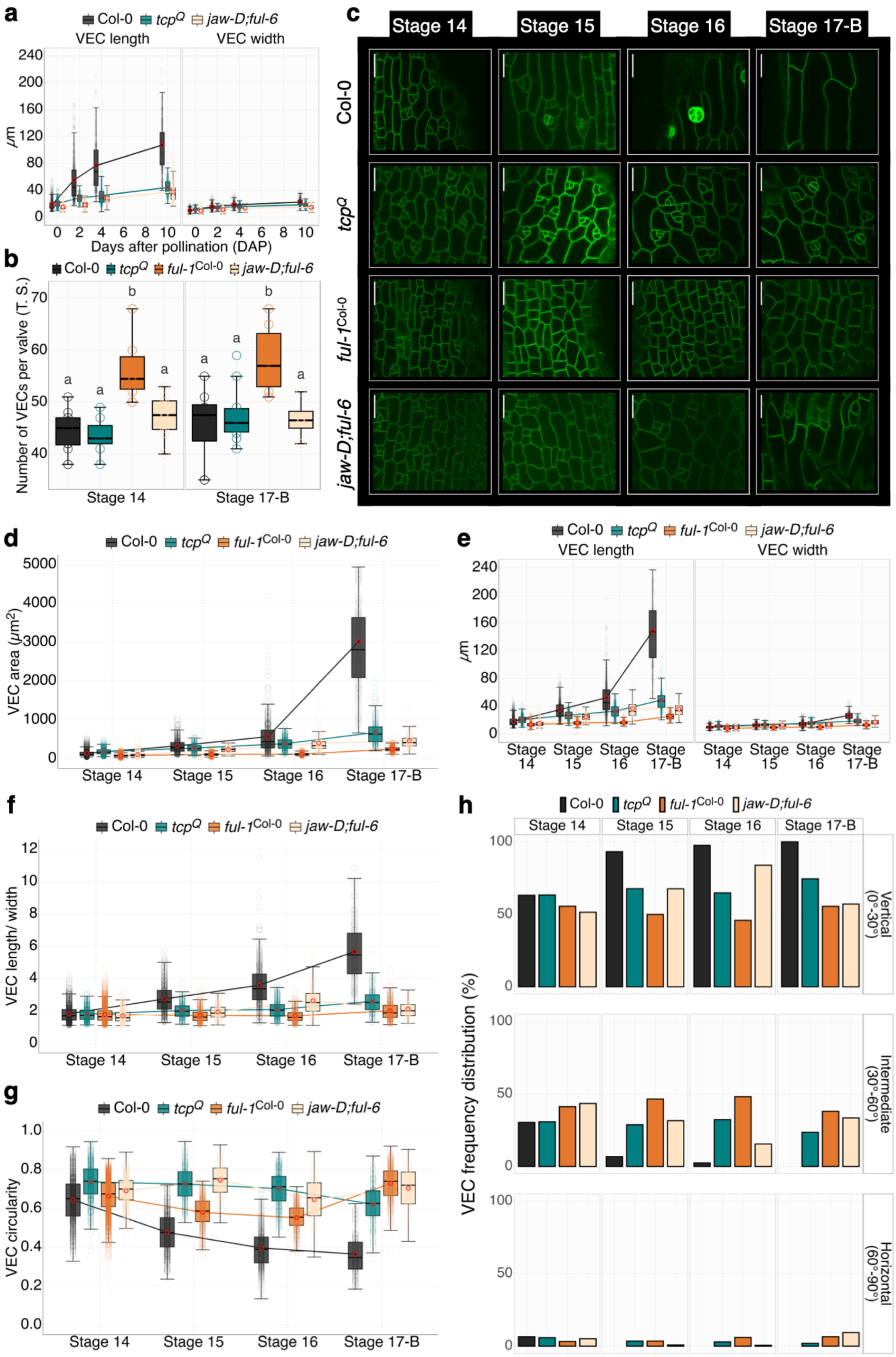


**Figure S4. Proximo-distal cell growth is reduced in the valve epidermal cells of *tcp* and**

***ful* mutants.**

(a) Length and width of the valve epidermal cells (VEC) at indicated days after pollination (DAP) along the X-axis. (b) Number of VECs (filled colored circles) in a transverse section of individual valves at indicated growth stages collected from 3 fruits of 3 individual plants. Variation in letters above the boxplots indicates statistical significance of $p<0.05$. One-way ANOVA followed by Tukey's HSD was used to determine significant differences. (c) Confocal fluorescence images of VECs in the indicated genotypes marked with *pUBQ10::ACYL-YFP* at indicated stages. Scale bar, 20 µm. (d-g) Area (d), length & width (e), length/width ratio (f) and circularity values (g) of the VECs of indicated genotypes at indicated growth stages. (h) Frequency distribution of the VECs analyzed in (d-g) among vertical, intermediate and horizontal categories in indicated genotypes at various stages of fruit growth. VECs were categorized as described in **Figure 4e**. VEC parameters are averages of cells quantified from 3-10 fruit valves from at least three independent plants across two independent batches, the individual means of which are indicated in open red circles. Colored circles indicate the individual cell parameter values. Each boxplot in (a-b, d-g) indicates the second and third quartile of data points, with the horizontal bar representing the median. The upper and the lower whiskers indicate either the highest/ lowest value or ±1.5 times the interquartile range.

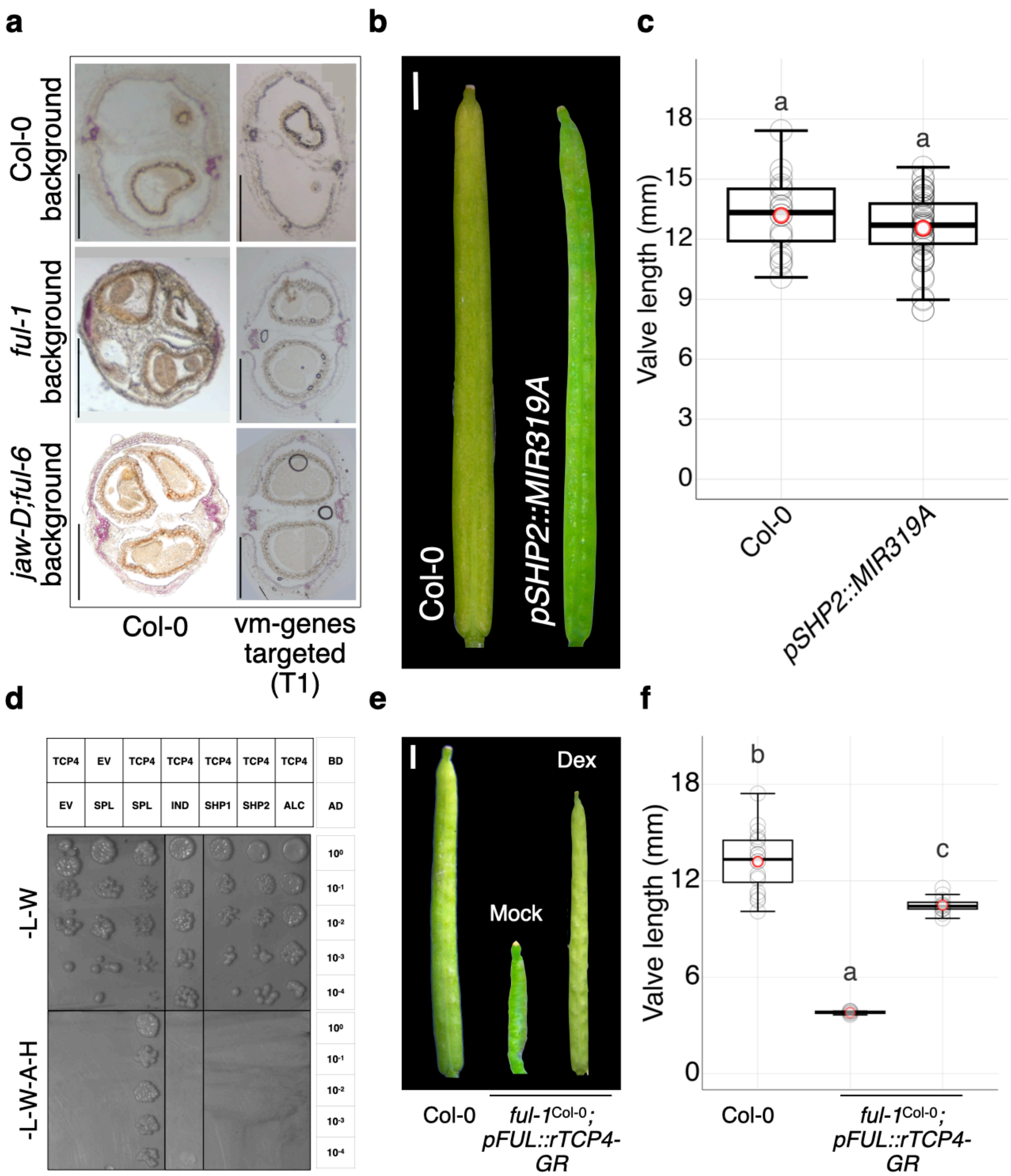


**Figure S5. Ectopic expression of valve margin genes and increased mesocarp lignification in the *jaw-tcp* mutants**

(a) Transverse sections of mature fruits of the indicated genotypes stained with phloroglucinol to reveal lignification (purple colour). 'WT control' refers to Col-0, and 'vm-genes targeted (T1)' refers to transgenic plants expressing a Cas9-editing construct targeting a single gRNA per gene to each of the four vm-genes. (b-c) Images of mature Col-0 and *pSHP2::MIR319A* fruits (b) and the valve length (c) of 20-50 fruits collected from at least three plants per grown batch across three independent batches. (d) Yeast two-hybrid assay to test physical interaction between TCP4-AD (top table, top row) and BD-domain fused with indicated proteins (top table, bottom row). Growth (at the indicated dilution, right table) in the -Leucine and -Tryptophan (-L-W, top panel) indicates the presence of both the AD and the BD constructs in the yeast cells, while growth in the -Leucine, -Tryptophan, -Adenine and -Histidine (-L-W-A-H, bottom panel) indicates physical interaction between the AD and BD construct expressed proteins. (e, f) Images of mature Col-0 and *ful-1;pFUL::rTCP4-GR* fruits (e) and their valve length (f) averaged from 11-19 fruits. Each boxplot in (c, f) indicates the second and third quartile of data points, with the horizontal bar representing the median. The upper and lower whiskers indicate either the highest & the lowest value or ±1.5 times the interquartile range. Filled grey and open red circles in (c, f) indicate the individual fruit valve length and average valve length, respectively. The difference in letters (marked above the boxplots in c, f) between genotypes indicates the statistical significance of $p<0.05$ as measured by an unpaired Student's *t*-test (c) and a one way Anova followed by Tukey's HSD (f). Scale bar, 200 μm (a), 1 mm (b, e).

### Supplementary table 1

| RT-qPCR | | |
|---|---|---|
| Primer number | Prime name | Sequence |
| P02631 | ARR16 FP | TTGCAAAGTGACAACAGCAGAG |
| P02632 | ARR16 RP | TCCAGGCATACAGTAATCGGTGA |
| P00109 | UBQ FP | GATCTTTGCCGGAAAACAATTGGAGGAGGATGGT |
| P00110 | UBQ RP | CGACTTGTCATTAGAAAGAAAGAGATACAGG |
| P06092 | FUL FP | CTCACGCGAAACCCAACAAA |
| P06093 | FUL RP | ACGACAACCCTGAAATGACCA |
| P06094 | SHP1 FP | GGAACTCAAAAACCTAGAAGGACG |
| P06095 | SHP1 RP | CTGCAAAATTCTCACAAAAGGGTG |
| P06096 | SHP2 FP | AAGGGTAAAAGAAATCGAGCTGC |
| P06097 | SHP2 RP | CCCCGACTGGTGAGAAGAAG |
| P06098 | IND FP | CTGGTGGTGCGAAGATGGA |
| P06099 | IND RP | ATCAGGGTTGGGAGTTGTGG |
| P06100 | ALC FP | CATTCCCAATTCCAACAAGACTGA |
| P06101 | ALC RP | GAGTTGGAGGTGGAACCTGT |
| P06761 | RPL FP | GCTGATTCCGATTTCCTCGC |
| P06762 | RPL RP | AGTCCGGCGTTAAAGGAATGA |
| ChIP-qPCR | | |
| Primer number | Prime name | Sequence |
| P03296 | ARR16 FP | AGATAAACCAAGCCATTCACTCTGGA |
| P03297 | ARR16 RP | GTCACAAATGGACCACCAATCTCTTT |
| P06656 | R1 FP | GAGTGTGGTCACACACAGAAAAA |
| P06657 | R1 RP | CGCACGTTGTCTTGTCAAACT |
| P06664 | R2 FP | ACGAAGCGGAAGAAATGGGA |
| P06665 | R2 FP | TTCTGTTAGGTGGGTTTTGATAACG |
| P06658 | R3 FP | ATGTTTAGGGCTCATCAACACAC |
| P06659 | R3 RP | AGGGTGTTCGGTTAGCAATTTTT |
| EMSA | | |
| Primer number | Prime name | Sequence |
| P06646 | BS1 forward | TCAATTCAACTAGGCCCAGTTTCCTCCTTC |
| P06647 | BS1 reverse | GAAGGAGGAAACTGGGCCTAGTTGAATTGA |
| P06648 | BS2 forward | GTGATGTTAAAATGGTCCTAAGTTGACATA |
| P06649 | BS2 reverse | TATGTCAACTTAGGACCATTTTAACATCAC |
| P06650 | BS3-4 forward | CAACACACAAATGTGGTCCATTTCTTTTTGTTCATGTGGTCCAATTTTGGTATAT |
| P06651 | BS3-4 reverse | ATATACCAAAATTGGACCACATGAACAAAAAGAAATGGACCACATTTGTGTGTTG |
| P06652 | BS5 forward | TTGCATATTGGTTGGTCCAAAAAGATCAAC |
| P06653 | BS5 reverse | GTTGATCTTTTTGGACCAACCAATATGCAA |
| Cloning | | |
| Primer number | Prime name | Sequence |
| P05033 | proFUL FP | GGGGACAACTTTGTATAGAAAAGTTGTACATCATCTGTATTTGTTT |
| P05034 | proFUL RP | GGGGACTGCTTTTTTGTACAAACTTGATCTCTCTCTCTTCAAAA |

| | | |
|---|---|---|
| P05035 | proSHP2 FP | GGGGACAACTTTGTATAGAAAAGTTGATCTCCAACGCATTGTTACG |
| P05036 | proSHP2 RP | GGGGACTGCTTTTTTGTACAAACTTGCATTTCTATAAGCCCTAGCTGAAG |
| P06190 | FUL cDNA FP | GGGGACAAGTTTGTACAAAAAAGCAGGCTATGGGAAGAGGTAGGGTTCAG |
| P06191 | FUL cDNA RP | GGGGACCACTTTGTACAAGAAAGCTGGGTCTCGTTCGTAGTGGTAGGACG |
| CRISPR/Cas9-editing | | |
| P06324 | FUL BS3-4 FP | ATTGTTGTTCATGTGGTCCAATTT |
| P06325 | FUL BS3-4 RP | AAACAAATTGGACCACATGAACAA |
| P06573 | SHP1 FP | ATTGGTCATATATAGGATCAATGG |
| P06574 | SHP1 RP | AAACCCATTGATCCTATATATGAC |
| P06575 | SHP2 FP | ATTGCTAGGGCTTATAGAAATGGA |
| P06576 | SHP2 RP | AAACTCCATTTCTATAAGCCCTAG |
| P06577 | IND FP | ATTGCACCATCTCCTCATGGATTG |
| P06578 | IND RP | AAACCAATCCATGAGGAGATGGTG |
| P06571 | ALC FP | ATTGGAGATGGGTGATTCTGACGT |
| P06572 | ALC RP | AAACACGTCAGAATCACCCATCTC |
| Genotyping | | |
| P00209 | TCP2 IS FP | TAGCATCTGAATTTCATAACCAATCTCGATACAC |
| P01269 | TCP2 IS RP | GGATTCTGCCGGTGATATCAAATGG |
| P01268 | TCP2 GS FP | CTCCTTCTTTAAATCCCAAACCAACC |
| P01269 | TCP2 GS RP | GGATTCTGCCGGTGATATCAAATGG |
| P03183 | TCP3 IS FP | AGAAACCTCCATGTGTGCTTG |
| P01996 | TCP3 IS RP | GACCTGCAGTTAATGGCGAGAATCGGATGAAGCA |
| P01994 | TCP3 GS FP | TTCGGATCCATGGCACCAGATAACGACCATTTCT |
| P01996 | TCP3 GS RP | GACCTGCAGTTAATGGCGAGAATCGGATGAAGCA |
| G5782 | TCP4 IS FP | ATGTCTGACGACCAATTCCATC |
| P00209 | TCP4 IS RP | TAGCATCTGAATTTCATAACCAATCTCGATACAC |
| G5782 | TCP4 GS FP | ATGTCTGACGACCAATTCCATC |
| G5783 | TCP4 GS RP | TCAATGGCGAGAAATAGAGGAA |
| P01088 | TCP10 IS FP | ACAAAGCAAGTGGGCAACAAAAACG |
| P02462 | TCP10 IS RP | TGGTTCACGTAGTGGGCCATCG |
| P01088 | TCP10 GS FP | ACAAAGCAAGTGGGCAACAAAAACG |
| P01089 | TCP10 GS RP | TAGTTTAGAGGTGTGAGTTTGGAGG |
| P02462 | TCP24 IS FP | TGGTTCACGTAGTGGGCCATCG |
| P01027 | TCP24 IS RP | TCCTTTCCTTTGCCTTGTCA |
| P04204 | TCP24 GS FP | GGGGACAAGTTTGTACAAAAAAGCAGGCTTCATGGAGGTTGACGAAGACATTGAG |
| P04205 | TCP24 GS RP | GGGGACCACTTTGTACAAGAAAGCTGGGTCCTATCTCCTTTCCTTTGCCTTGTC |
| P05715 | *stk-2* FP | GCTTGTTCTGATAGCACCAACACTAGCA |
| P05705 | *stk-2* RP | GGGGACCACTTTGTACAAGAAAGCTGGGTGGTCATATGAATCACTAAATAAGATAC |
| P05804 | *ful-6* FP | GTTTGCAACCAATATCAAATCACAA |
| P05805 | *ful-6* RP | ATGGTGAGATGACATACTGTAGTT |

| | | |
|---|---|---|
| P06736 | SHP1 FP | TGAGAACTCAGGTAAGGTTGTGAA |
| P06737 | SHP1 RP | GCAACTTCGGCATCACACAA |
| P06738 | SHP2 FP | GCTAAGCCAGGTATGGTTATTGATG |
| P06739 | SHP2 RP | TGCGTCGTTTGCAGAAAGTG |
| P06740 | ALC FP | CTGTTGCGAGACCTTACTTTTCA |
| P06830 | ALC RP | GAGACTCCATAACCGACGGA |
| P06742 | IND FP | AGTGTGCGACTCTTGTGTCT |
| P06743 | IND RP | ACTGCATCTCCTTCATCGCA |
| P06520 | FUL FP | CATGCGGTCTTGAGATGTGG |
| P06521 | FUL RP | GACAAACACTCGTCCGACTAA |
